# The orphan histidine kinase TodK controls *Myxococcus xanthus* biofilm development by inactivating the CRP/Fnr homolog, MrpC

**DOI:** 10.64898/2026.07.28.741289

**Authors:** Christopher Mataczynski, Maike Glaser, Stuart Huntley, Penelope I. Higgs

## Abstract

Environmental bacteria have abundant signaling systems wired into complex gene regulatory networks to adapt to fluctuating conditions. In *Myxococcus xanthus*, starvation triggers a developmental program (specialized biofilm) that produces spore-filled multicellular fruiting bodies surrounded by a distinct quiescent state termed peripheral rods. Fruiting body structure as well as the proportion of cells following each fate can be tuned by a large repertoire of signaling proteins, including numerous orphan histidine kinases. Here, we focus on the histidine kinase TodK which was previously demonstrated to influence developmental progression. We find that loss of TodK produces distinct developmental phenotypes that vary with environmental conditions. To quantify these effects, we developed an image-analysis pipeline that measures aggregation and fruiting body patterning during development on nutrient-limited agar. These analyses revealed the *todK* mutant precociously aggregates particularly at the peripheries of the colony. Under submerged-culture conditions, initial production of aggregates was not accelerated but aggregates exhibited accelerated progression to mature fruiting bodies. Overexpression of active TodK completely blocked fruiting body formation. Molecular analyses demonstrated that TodK overproduction suppressed expression of core developmental regulators including FruA and CsgA (C-signal). Interestingly, protein accumulation of MrpC, necessary for expression of both FruA and the C-signal was not significantly perturbed suggesting TodK silences MrpC transcriptional activity. Together, these findings establish TodK as a modulator of developmental progression and demonstrate how quantitative phenotyping approaches can reveal biologically meaningful functions for orphan histidine kinases whose mutant phenotypes might otherwise appear subtle.

**Summary Statement:** Quantitative analysis of multicellular development reveals previously hidden functions of an orphan histidine kinase, highlighting the importance of robust phenotyping approaches for understanding bacterial signaling networks.

## Introduction

To survive in fluctuating environments, free-living bacteria have evolved remarkably complex developmental and multicellular behaviors, including biofilm formation, cellular differentiation, and phenotypic heterogeneity (Chong and Shapiro, 2024). These processes are coordinated by regulatory networks that integrate environmental information and direct large-scale changes in gene expression. A central challenge in understanding such systems is identifying how sensory proteins modulate developmental decisions in response to changing conditions. In many cases, disruption of individual signaling proteins produces only subtle phenotypic effects, reflecting extensive network redundancy and buffering. Consequently, sensitive quantitative approaches are often required to uncover the biological functions of regulatory proteins and to understand how environmental inputs shape developmental outcomes.

The social bacterium *Myxococcus xanthus* provides a powerful model for investigating environmentally responsive developmental regulation (Kroos *et al*., 2025). Starvation triggers a multicellular developmental program (a specialized biofilm) in which some cells first aggregate into hay-stacked shaped mounds of ∼10^5^ cells and subsequently differentiate into resistant spores to produce mature fruiting bodies. Other cells follow different cell fates: differentiation into a distinct quiescent state arrayed around the fruiting bodies (peripheral rods) (O’Connor and Zusman, 1991a; Higgs *et al*., 2014), or cell lysis (Wireman and Dworkin, 1977; Janssen and Dworkin, 1985; Lee *et al*., 2012). The proportion of cells in each fate and the rate at which development progresses is tuned to different environmental conditions (O’Connor and Zusman, 1991b; Marcos-Torres *et al*., 2020; Kasto and Higgs, 2025). Because developmental outcomes can be readily observed at the population level, *M. xanthus* has long served as a model system for linking signaling pathways to multicellular behavior (Kroos *et al*., 2025).

The *M. xanthus* developmental program is directed by a core signal-responsive genetic regulatory network (Kroos, 2017), but the mechanisms by which the program can be tuned to environmental conditions are not well elucidated. The developmental program is initiated by starvation via the stringent response, and can proceed by when a subsequent population density sensing requirement (A-signaling) is met (Harris *et al*., 1998). At the core of the developmental program are three unusual regulatory factors: the transcription factors MrpC and FruA, and the C-signal derived from the CsgA protein (Shimkets and Rafiee, 1990; Ogawa *et al*., 1996; Sun and Shi, 2001a). MrpC, a member of the CRP/FNR transcription factor superfamily, lies at the top of the regulatory hierarchy. MrpC activity does not appear to be regulated by an endogenous ligand (Ueki and Inouye, 2003; McLaughlin *et al*., 2018; Saha *et al*., 2019b; Feeley *et al*., 2019). Instead, MrpC accumulation may be key to its regulatory role, with different MrpC accumulation thresholds associated with distinct developmental patterns (Lee, 2009; Schramm *et al*., 2012; Marcos-Torres *et al*., 2020; Lall *et al*., 2024). Consistently, MrpC is subject to multiple levels of regulation. *mrpC* transcription is induced by the enhancer binding protein, MrpB (Sun and Shi, 2001b), controlled by the AskABC histidine kinases (HKs) (Marcos-Torres *et al*., 2020), and by negative autoregulation (McLaughlin et al., 2018). Post-translational regulation occurs through regulated proteolysis and by Ser/Thr kinase-dependent phosphorylation. Regulated proteolysis can be triggered by the EspAC HKs, or by return of nutrients (Higgs *et al*., 2008; Schramm *et al*., 2012; Rajagopalan and Kroos, 2014; Hoang and Kroos, 2018; Kasto and Higgs, 2025). Post-translational modification occurs by the Ser/Thr kinase, Pkn14 (Nariya and Inouye, 2005; Feeley *et al*., 2019).

Initial MrpC accumulation induces expression of FruA, an atypical response regulator that lacks residues necessary for phosphorylation in its receiver domain (Ellehauge *et al*., 1998; Ueki and Inouye, 2003). FruA activates and/or represses hundreds of genes, some in cooperation with MrpC and/or dependent on its activation state (Mittal and Kroos, 2009; McLoon *et al*., 2021; Farrugia *et al*., 2025). Activation of FruA is somehow dependent upon the C-signal, proposed to be a cell-cell contact morphogen (Ellehauge *et al*., 1998). C-signal is derived from the *csgA* gene which encodes a 25 kDa protein that shares homology to short chain alcohol dehydrogenases. *csgA* mutants do not aggregate nor sporulate, but exogenous C-signal supplied as a purified protein or by co-development with wild type cells, can restore fruiting body formation. Low levels of C-signal are proposed to somehow activate FruA to induce aggregation, which, in turn, increases cell-cell contact as the cells aggregate into mounds (Kim and Kaiser, 1990; Kim and Kaiser, 1991; Li *et al*., 1992; Kruse *et al*., 2001; Lobedanz and Sogaard-Andersen, 2003; Hoang *et al*., 2021). Inside mounds, intense cell-cell contact leads to highly active FruA which is required to induce transcription of sporulation genes (Ellehauge *et al*., 1998; Saha *et al*., 2019a; Hoang *et al*., 2024).

In addition to the core developmental regulatory network, numerous HKs influence progression through development. Histidine kinases are particularly abundant in environmental bacteria and often function within multi-component signaling networks that integrate diverse environmental cues (Galperin, 2005; Francis and Porter, 2019). In *M. xanthus*, many kinase genes have been implicated in regulating developmental progression (Cho and Zusman, 1999; Rasmussen and Sogaard-Andersen, 2003; Lee *et al*., 2005; Higgs *et al*., 2005; Higgs *et al*., 2008; Shi *et al*., 2008; Jagadeesan *et al*., 2009; Glaser and Higgs, 2019), yet relatively few have been characterized mechanistically. A recurring challenge is that deletion mutants often exhibit subtle developmental defects that can be overlooked using qualitative approaches. Consequently, systematic and quantitative characterization of developmental phenotypes is required to reveal the contributions of these signaling proteins to developmental regulation.

TodK is a predicted cytoplasmic HK that has previously been hypothesized to regulate the timing of *M. xanthus* fruiting body formation by delaying C-signal-dependent activation of FruA (Rasmussen and Sogaard-Andersen, 2003). As TodK is genetically orphan and the stimuli it is responding to are unknown, exactly how it modulates C-signal reception by FruA is unclear. Further characterization of TodK therefore represents an attractive model for deeper understanding of how core developmental regulatory networks can be modulated by HKs in response to environmental conditions. In this study, we capitalize on the observation that a Δ*todK* mutant produces subtly distinct phenotypes on nutrient limited agar (NLA) vs submerged culture (SC) developmental conditions. We developed an image-analysis workflow to measure aggregation and spatial fruiting body patterning during development on NLA and applied this approach to identify more precise developmental defects associated with loss of TodK. Using both loss- and gain-of-function analyses, we demonstrate that TodK appears to silence MrpC-dependent activation of targets including *fruA*, *csgA*, and *espA*. We propose a model in which local nutrient levels influence TodK to control MrpC-dependent induction of developmental transitions. Our findings illustrate how quantitative phenotyping can uncover functional roles for signaling proteins whose contributions to development may otherwise remain hidden.

## Results

### TodK controls onset of developmental transitions in response to environmental conditions

TodK is a cytoplasmic histidine kinase homolog which contains amino terminal tandem PAS sensing domains (Rasmussen and Sogaard-Andersen, 2003)(Fig. 1A). To extend on previous work characterizing TodK, we generated a *todK* in-frame deletion (Δ*todK*) in the alternate *M. xanthus* wild-type (WT) strain, DZ2 and assessed the developmental phenotypes on Clone Fruiting (CF) nutrient-limited agar (NLA) plates (Fig. 1B) and under submerged culture (SC) (Fig. 1F and Movies S1 and S2). By 48 hours of development on NLA, the two strains produced distinct mature fruiting body patterns: while the wild type produced typical fruiting bodies throughout the area of the initial spotted cells, the Δ*todK* mutant produced smaller and more numerous fruiting bodies throughout the spot with a ring of tightly clustered fruiting bodies at the spot periphery (Fig. 1B).

**Fig. 1.**
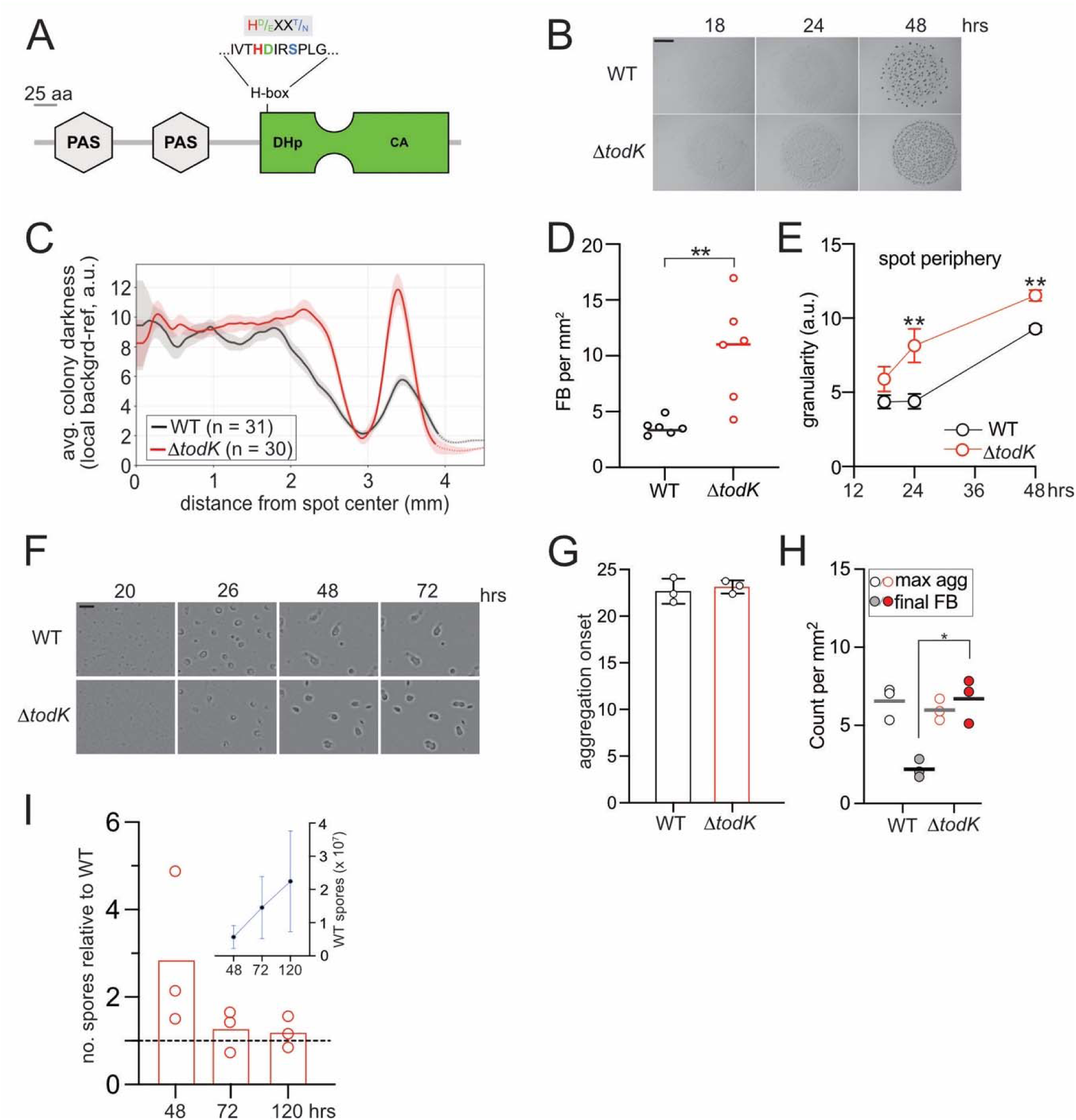
TodK influences fruiting body patterning and dynamics in response to the distinct environmental conditions. Strains were induced to develop on nutrient-limited CF agar plates (**B-E**) or in submerged culture (**F-I**). **(A)** TodK domain architecture. PAS, Per-ARNT-Sim sensing domain (Möglich *et al*., 2009); green shape: histidine kinase region containing dimerization / phosphoaccepting (DHp) and catalytic ATPase (CA) domains; H-box, sequence containing the phosphorylated histidine (H; red) and residues predicted to govern kinase (green) and/or phosphatase (blue) activity. Scale bar: 25 residues. **(B)** Developmental time course of 10 μl (at 4 x10^9^ cells ml^-1^) of wild-type (WT; DZ2) or Δ*todK* (PH1332) cells spotted on CF agar plates and imaged at the indicated hours. Scale bar: 2 mm. **(C)** Radial fruiting body patterns produced at 48 h. Mean radial darkness in arbitrary units (a.u.) detected in wild type (black) or Δ*todK* (red) developmental spots plotted against absolute distance from the spot center in mm. Solid line, full 360-degree coverage; Dotted line, partial coverage; Shaded band, standard error of the mean. Indicated n values are from six independent biological replicates each containing ≥ 3 technical replicates. **(D)** Average fruiting bodies per mm^2^ at 48 h development. Fruiting bodies were detected as dark objects on a local-background-referenced darkness map. Circles are the average of ≥ 3 technical replicates per biological replicate. Lines are the median from n = 6 independent biological replicates. Data presented is graphical representation of that in Table 1. **(E)** Aggregation analysis at the spot periphery region. Image texture (granularity) in a.u. averaged over the spot periphery was measured for cells developing at 18, 24 and 48 hours. Values represent the mean ± SEM (n=6 independent biological experiments). Technical replicates (≥ 3) per experiment) were averaged to determine the value for each biological replicate. **(F)** Developmental time course of wild type and Δ*todK* strains developing under submerged culture (SC) in 96 well plates. Representative images are shown from a time course in which cells were imaged every 30 min for 72 hours. Scale bar, 0.25 mm. **(G)** Aggregation onset calculated as the average hour of development at which visible aggregates are first observed. Values represent the mean ± SEM (n = 3 independent biological experiments). Technical replicates (≥3) per experiment) were averaged to determine the value for each biological replicate. **(H)** Aggregate and fruiting body counts per mm^2^ in SC. Maximum initial aggregates (open circles) and final fruiting bodies observed at 72 h of development (filled circles). Data points represent the average of ≥ 3 technical replicates and the mean (bar) is the average of three independent biological replicates. **(I)** Sporulation efficiency of the Δ*todK* relative to the wild type at the indicated hours. Wild type and Δ*todK* strains were induced to develop by submerged culture in 24-well tissue culture plates. At the indicated times, heat and sonication resistant spores enumerated from the Δ*todK* strain were normalized to the wild type. Bars represent the mean of three independent biological replicates; circles are the average of three technical replicates. Dashed line, normalization line. Inset: Average number of spores per well produced by the wild type over time. Values are the mean and associated standard deviation from 3 independent biological replicates. Statistical significance was tested by Mann-Whitney U **(D)**, Welch’s t-test **(G, H, I)** or paired t-test **(E)**. Statistically significant differences are indicated as follows: *, p ≤ 0.05; **, p ≤ 0.01.

**Table 1.** Fruiting body spatial analysis on strains developing for 48 hours on CF agar.

| Metric (per spot) | WT | *ΔtodK* | p (6 vs 6)<sup>a</sup> |
| --- | --- | --- | --- |
| Fruiting-body count | 158 ± 46 | 383 ± 166 | 0.004 |
| FB density (per mm<sup>2</sup>) | 3.3 ± 1.0 | 8.7 ± 4.3 | 0.004 |
| Mean nearest-neighbor distance (NND) (mm) | 0.266 ± 0.051 | 0.182 ± 0.032 | 0.002 |

<sup>a</sup> probability of significance as assessed by a two-sided Mann–Whitney U on six per-experiment means.

To quantify this phenomenon, we first developed a tool to measure the average radial darkness on images as absolute pixel intensity from the center of the spotted cells to the frame edges at 48 hours of development and then averaged these densities from all images from multiple biological replicates. As a proof of concept, we first compared the developmental spots from wild type vs the non-developing Δ*mrpC* strain in paired experiments containing at least 8 images from four biological replicates (Fig. S1 A and B). This analysis demonstrated that the wild type reached an average pixel density of ∼12 arbitrary units (a.u.) at the spot center (0 mm), while the Δ*mrpC* strain produced a consistently low level of colony darkness (∼2.5 a.u.) from 0 to ∼2.7 mm radius (Fig. S1 A, B). Both strains produced a small peak near the original spot periphery (∼ 5 a.u. at 3.6 mm for the wild type vs ∼ 3.8 a.u. at 3 mm for Δ*mrpC* strains). We next compared the wild type and Δ*todK* strains using at least 30 images from six independent biological replicates. The wild type again exhibited a high pixel density at spot center that gradually decreased to a low-density zone at 3 mm followed by the small edge peak (Fig. 1C). In contrast, the Δ*todK* strain maintained the high pixel density from 0 - 2.5 mm followed by a more sudden drop into low density zone (∼ 3 mm) and then a much sharper peak (∼12 a.u.) at the spot periphery before dropping to background levels (Fig. 1C). These results provide quantitative evidence that the Δ*todK* mutant consistently produces a distinct fruiting body patterning in which the core of the original spot is filled with numerous fruiting bodies, surrounded by a body-poor ring and then a dense band of tightly clustered fruiting bodies near the spot periphery.

In addition to displaying a novel fruiting body patterning, we developed a tool to quantify spatial metric of fruiting bodies from 48 hours of development (Fig. S1C). These analyses indicated that relative to wild type, the Δ*todK* mutant produced 2.4-fold more fruiting bodies per spot at 2.6-fold higher density (Table 1 and Fig. 1D). Consistently, the Δ*todK* fruiting bodies are ∼30% more closely packed together as measured by average nearest neighbor distance (NND) (Table 1). The observation that the Δ*todK* strain produced more numerous fruiting bodies was noted previously (Rasmussen and Sogaard-Andersen, 2003).

It was also previously observed that a Δ*todK* mutant begins to aggregate earlier than the wild type (Rasmussen and Sogaard-Andersen, 2003). This phenotype was not consistently obvious, because initial aggregates were sometimes poorly defined in the spot center such that detection of early aggregation was difficult. Additionally, because these are static images and aggregation onset can fluctuate slightly between biological replicates, we sought a quantitative method to define small differences in aggregation relative to the wild type in each biological replicate. For this purpose, we developed a method to assess spot texture by measuring granularity of the spotted cells in the core and periphery of the spotted cells from 18, 24, and 48 hr images (Fig. S1D). These analyses demonstrated that mean Δ*todK* granularity trended higher than the wild type in the 24 and 48 hr time points (Fig. S1E). However, it was most pronounced at the spot periphery, showing a statistically significant increase between 18 and 24 hours, while the wild type remained relatively constant (Fig. 1E). This observation is consistent with large peak in radial density at the spot periphery (Fig. 1B and C). Together, these results hint that in wild-type cells, TodK is most active at the cell periphery. After at least 18 hrs after spotting, some cells migrate into the surrounding area, likely reducing the local cell density at the periphery of the spot. TodK may be responding either to the locally reduced cell density or to increased nutrient availability at the spot periphery, where nutrient consumption is lower simply because fewer cells occupy the area.

We next analyzed wild type and Δ*todK* development by submerged culture (SC) in which cells are induced to develop in tissue culture wells, producing a uniform cell density per unit area. Furthermore, SC enables automated image capture in a plate reader allowing more detailed analyses of developmental dynamics (Glaser and Higgs, 2019)(Fig. 1F and Movies S1 and S2). Interestingly, under these developmental conditions, the WT and Δ*todK* strains exhibited no significant difference in early developmental events: both the onset of visible aggregation centers (23 ± 1 and 23.1 ± 0.7 hours, respectively) (Fig. 1G), and the maximum number of initial aggregation centers (6.6 ± 1.1 and 6.0 ± 0.7 per mm^2^, respectively) (Fig. 1H) were not significantly different. However, later developmental events differed between the WT and Δ*todK* strains. In the WT strain, 67 ± 4% of the initial aggregates dissolved (Fig. 1H) and the remaining aggregates were mobile for 31 ± 7 h (Movie S1). By 50 h, all remaining WT aggregates had transitioned into darkened and immobile mature fruiting bodies (FBs) (Fig. 1H, Movie S1). In contrast, none of the Δ*todK* initial aggregates dissolved and no significant aggregate mobility was observed (Fig. 1H, Movie S2). Thus, mature FBs were produced 26 ± 6 hours earlier than in the wild type, and the final number of FBs was similar to the number of initial aggregates (Fig. 1H).

We speculated the differences in aggregate kinetics could arise if TodK affects the onset of sporulation, because myxospores are immobile. To investigate this hypothesis, heat- and sonication-resistant spores were harvested and enumerated from cells developing under submerged culture in 24 well plates at 48, 72, and 120 h of development. The Δ*todK* strain reproducibly produced more spores relative to the WT at 48 h (although it did not meet statistical significance criteria) but similar sporulation levels to the WT 72 and 120 hours (Fig. 1I). Together, these results suggest that under submerged culture, TodK does not appear to affect aggregation in wild-type cells but likely delays sporulation, thereby delaying the transition of aggregates into mature FBs. Unlike cells on agar, cells in tissue culture plates do not experience edge-associated gradients in cell density and likely experience more uniform nutrient availability.

### Over-production of active TodK prevents development

Constitutive and/or overexpression of regulatory proteins can often result in magnification of phenotypes. To gain further clarity on the role of TodK during development, we cloned *todK* behind the highly active pilin promoter (P_pilA_) (Wu and Kaiser, 1997), and integrated the construct into the Δ*todK* background at the Mx8 phage attachment site (Δ*todK attB*::P_pilA_-*todK*; hereafter *todK*^++^). Strikingly, when analyzed for development under SC, the *todK*^++^ strain failed to begin aggregation and produced no heat- and sonication-resistant spores (Fig. 2 and Movie S3). To determine if TodK activity was necessary for this phenotype, we also generated a similar construct in which the predicted site of autophosphorylation, the histidine residue at position 275 (H_275_), was substituted with an alanine (*todK*^++^) (Fig. 1A). The *todK* ^++^ strain produced aggregates, mature FBs, and sporulation levels with similar characteristics to the Δ*todK* strain (Fig. 2 and Movie S4). We confirmed that TodK levels were stable and overexpressed in the *todK*^++^ and *todK_H275A_*^++^ strains by anti-TodK immunoblot analysis of cell lysates harvested at 0, 18, 24 and 30 h of development. In wild-type cell lysates, a band corresponding to the predicted molecular mass of TodK (55 kDa) was barely detected from 0-24 hours and more apparent at 30 hours, suggesting it was produced at very low levels (Fig. 2B). However, in the *todK*^++^ and *todK*++ _H275A_ backgrounds, the ∼55 kDa band was readily detected at all timepoints (Fig. 2B). As confirmation, mass spectrometry analysis of WT, *todK*^++^, and *todK*^++^ lysates at 30 hours PS indicated that TodK and TodK_H275A_ were ∼100-fold more abundant in the overexpression strains (data not shown). Together, these results suggested that overexpression of TodK inhibits development, and that the kinase activity was necessary for this inhibition.

**Fig. 2.**
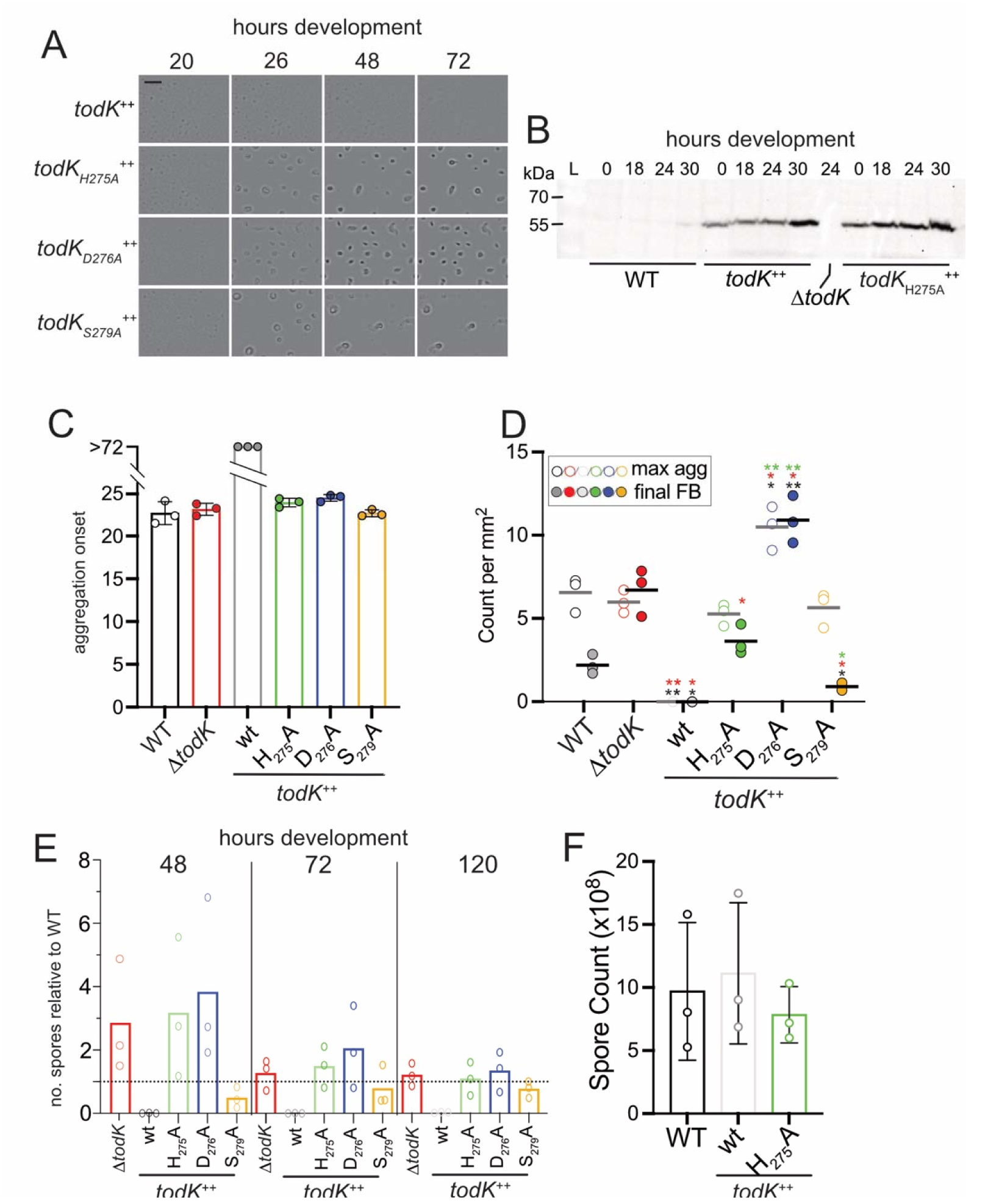
Overexpression of active TodK prevents development. **(A)** Submerged culture developmental phenotypes of strains overexpressing wild type, predicted kinase (K)- and/or predicted phosphatase (P)-negative *todK* alleles from strains PH2026 (Δ*todK attB*::P_pilA_-*todK*; *todK*^++^), PH2027 (*todK_H275A_* ^++^; K^-^ P^-^), PH2028 (*todK _D276A_*^++^; K^-^P^+^) or PH2029 (*todK_S279A_*^++^; K^+^P^-^) in a 96-well microtiter plate. Strains were imagined every 30 minutes from 0 to 72 h; images from the indicated hours are displayed. Scale bar: 250 μm. **(B)** anti-TodK immunoblot analysis of protein lysates prepared from wild type (WT; DZ2), *todK*^++^, or *todK* ^++^, strains developed for 0-30 hr. Antibody specificity control is lysate from Δ*todK* (PH1048) at 24 hour development. **(C,D)** Aggregation and fruiting body dynamics of the strains filmed in **A**. **(C)** Aggregation onset calculated as the average hour of development at which visible aggregates are first observed. Values represent the mean ± SEM (n=3 independent biological experiments). Technical replicates (≥3) per experiment) were averaged to determine the value for each biological replicate. **(D)** Aggregate and fruiting body counts per mm^2^ in the indicated strains. Maximum initial aggregates (open circles) and final fruiting bodies observed at 72 h of development (filled circles). Data points represent the average of ≥3 technical replicates and the mean (bar) is the average of three independent biological replicates. Colored stars represent statistical significance difference in the maximum aggregates or final fruiting bodies relative to the corresponding strain color. **(E)** Sporulation efficiency of the indicated strains relative to the wild type at the indicated hours. Strains were induced to develop by submerged culture in 24-well tissue culture plates. At the indicated times, heat and sonication resistant spores enumerated from each strain were normalized to the wild type. Bars represent the mean of three independent biological replicates; circles are the average of three technical replicates. Dashed line, normalization line. **(F)** Heat and sonication-resistant chemically-induced spores harvested vegetative broth cultures treated with 0.5 M glycerol for 24 hours. Bars and whiskers represent the means and associated SD from three independent biological replicates. **(C-F)** Statistical significance was tested by Welch’s t-tests. Statistically significant differences are indicated as follows: *, p ≤ 0.05; **, p ≤ 0.01. The WT and Δ*todK* data in panels **C-E** is the same data as in Fig. 1 and added here for reference.

During the developmental program, cells are only induced to sporulate after aggregation into mounds. To clarify whether the sporulation defect observed in the *todK*^++^ mutant was specifically due to disruption of the core sporulation program, we assayed WT, *todK*^++^, and *todK*++ _H275A_ chemical induction of sporulation in vegetative broth cultures, a method that bypasses the requirement for aggregation (Dworkin and Gibson, 1964). All strains produced similar levels of chemically induced spores (Fig. 2F). This result suggests that overexpression of active TodK affects processes necessary to induce aggregation in the genetic regulatory network, and that failure to sporulate in submerged culture is a result of the lack of production of mounds, rather than a defect in the core sporulation program.

### TodK does not autophosphorylate *in vitro*

To examine if TodK could undergo autophosphorylation *in vitro*, we purified Thioredoxin-histidine tagged TodK (Trx-His_6_-TodK_wt_) and the putative kinase inactive form (Trx-His_6_-TodK_H275A_). Purified proteins were incubated with radioactively labeled [γ^32^P]-ATP under standard conditions with reaction buffer containing 5 mM MgCl_2_. Surprisingly, no *in vitro* autophosphorylation was detected even after 60 minutes of incubation with Trx-His_6_-TodK_wt_ (Fig. S3A), whereas concomitant autophosphorylation of a control histidine kinase, Trx-His_6_-SinK (Glaser and Higgs, 2019), was readily detectable (Fig. S3B). Neither increasing the MgCl_2_ concentration to 10 mM nor using alternate cations of 1 or 10 mM MnCl_2_ or CaCl_2_, which have previous been shown to induce kinase activity in HKs (Gamble et al., 1998; Psakis et al., 2011), promoted detectable on autophosphorylation (Fig. S3C). To examine whether the putative PAS sensing domains inhibited autophosphorylation, we purified the kinase region lacking the putative sensor region (Trx-His_6_-TodK_kinase_) and PAS (Trx-His_6_-TodK_PAS_) domains. Autophosphorylation was not observed if the TodK_kinase_ was incubated alone nor if the TodK_kinase_ and TodK_PAS_ regions were combined in equimolar ratio, even after 45 minutes of incubation (data not shown).

### Genetic analysis suggests TodK^++^ kinase and phosphatase activities are essential to inhibit development

Most HKs are bifunctional, displaying both kinase and phosphatase activity on targets (Stock et al., 2000). Since we were unable to detect TodK autophosphorylation *in vitro*, we hypothesized that TodK may be a dedicated phosphatase *in vivo*. Consistently, it has previously been reported that substitutions of the invariant histidine residue in HKs disable phosphatase activity as well as kinase activity (Huynh et al., 2010; Willett and Kirby, 2012). Importantly, other residues in the H-box motif (Fig. 1A) are essential and specific for kinase or phosphatase activity separately. Specifically, substitution of the Asp or Glu (D/E) reside located immediately downstream of the invariant histidine; H +1) renders HKs kinase dead but phosphatase active (K-P+). In contrast, substitution of the Thr or Asn (T/N) residue located at the H +4 position renders the HK kinase-active but phosphatase-dead (K+P-) (Willett and Kirby, 2012). TodK contains a functionally conserved Ser (S) residue at the H + 4 position (Fig. 1A). Thus, we generated alanine substitutions of D_276_ or S_279_ in *todK*^++^ to generate *todK_D276A_*^++^ or *todK*_S279A_^++^ strains (putatively K^-^ P^+^, and K^+^ P^-^, respectively). All three TodK^++^ H-box mutant strains produced similar levels of TodK protein (data not shown). Surprisingly, both predicted K-P+ and K+P-TodK^++^ mutants phenocopied the *todK_H275A_*^++^ (K^-^ P^-^) strain by restoring development with aggregation onset indistinguishable from WT (Fig. 2A, C). However, subtle differences in later developmental phenotypes could be detected (Movies S5 and S6). The *todK_D276A_*^++^ K^-^ P^+^ strain produced 1.6-fold more aggregation centers than observed in any of the other mutants and these aggregation centers failed to dissolve (Fig. 2D). Sporulation levels at 48 hours were consistently higher than the WT, although they did not reach criteria for statistical significance (Fig. 2E). In contrast, the *todK*_S279A_ K^+^ P^-^ strain developed most similarly to WT, with aggregates that dissolved into fewer mature FBs and comparable sporulation levels at 48 h (Fig. 2C, E).

Generation of the H_275_A, D_276_A, and S_279_A substitution mutations in the native genomic *todK* locus produced the average distinct fruiting body patterning and early aggregation phenotypes that resembled the Δ*todK* mutant when analyzed on nutrient limited CF agar plates (Fig. S2).Together, these results suggest that both kinase and phosphatase activities are necessary for overexpressed TodK to inhibit the developmental program, or for native TodK to appropriately control aggregation and fruiting body patterning on CF agar. However, as we could not detect TodK autophosphorylation *in vitro* and the target(s) of TodK kinase/phosphatase activity is unknown, we could not biochemically verify that these mutations specifically abolish kinase and/or phosphatase activity as predicted.

### C-signaling is abolished by overexpressing TodK

We next turned our attention to the mechanism by which TodK^++^ inhibits development. TodK was previously suggested to act on the C-signal transduction pathway (Rasmussen and Sogaard-Andersen, 2003). Since the *todK*^++^ and *csgA* developmental phenotypes are similar (*i.e.* both strains failed to develop), we next examined CsgA levels in the *todK*^++^ strain. Anti-CsgA immunoblot analysis on cell lysates harvested from WT, or *todK*^++^ strains developing between 0 and 30 h, revealed that in the WT strain, the expected 25 kDa CsgA band was produced after the onset of starvation and accumulated by 18 h; this band was absent in a *csgA* mutant (Fig. 3A). In the *todK*^++^ strain, CsgA levels were strikingly reduced at all time-points examined. Furthermore, qRT-PCR analysis on RNA harvested from cells at 0, 6, and 24 h PS (Fig. 3B) indicated at least a 2-fold decrease in *csgA* mRNA levels compared to WT. These results suggest that overexpression of TodK blocks development by preventing appropriate *csgA* expression.

**Fig. 3.**
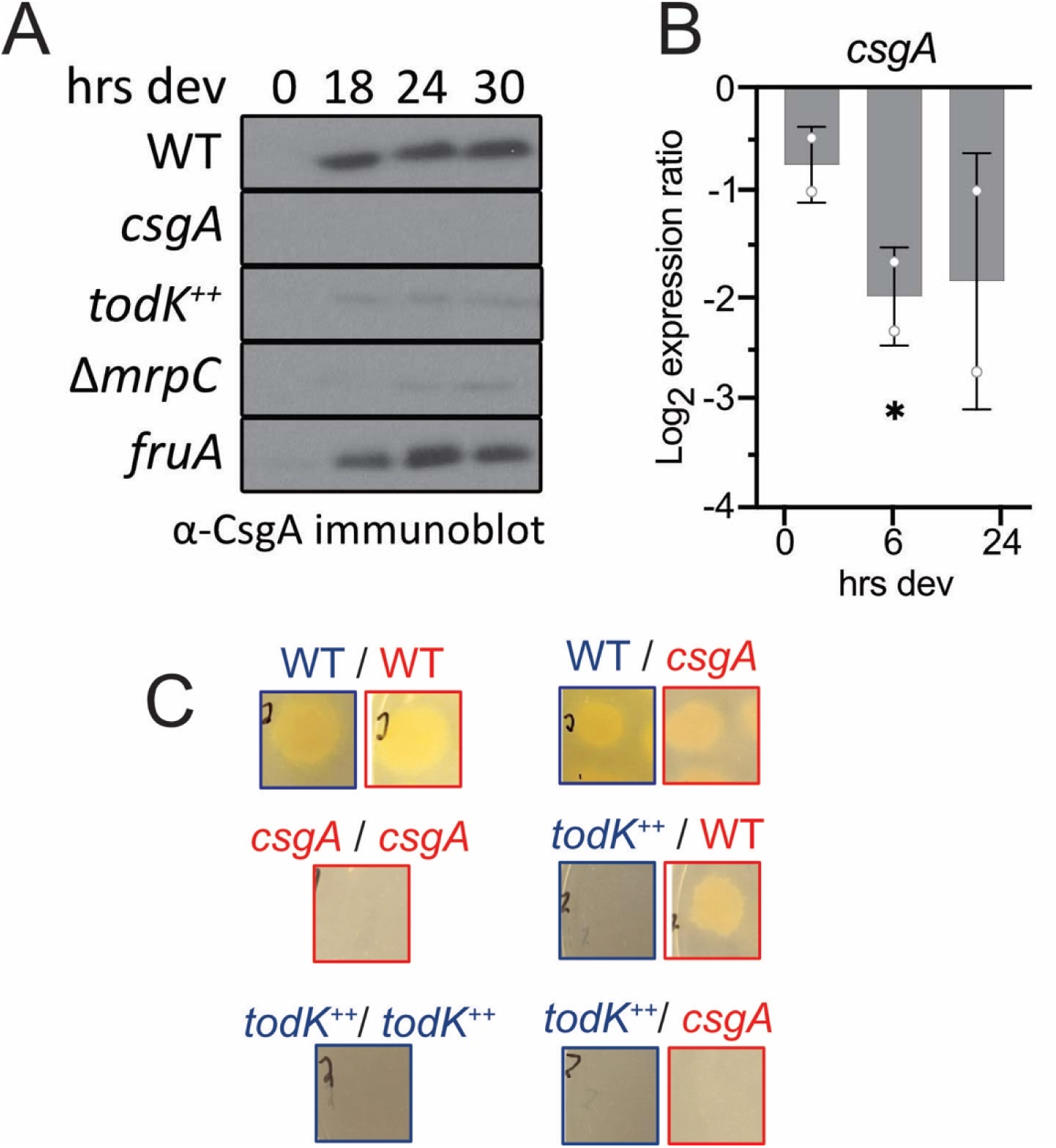
Overexpression of TodK prevents CsgA accumulation. **(A)** anti-CsgA immunoblot analysis of WT (DZ2), *csgA*::Tn5 (PH1014), *todK*^++^ (PH2030), Δ*mrpC* (PH1025), and *fruA::pPH128* (PH1013) strains developing under submerged culture conditions at indicated hours. **(B)** qRT-PCR analysis of *csgA* mRNA in the *todK*^++^ strain. RNA was harvested from cells developed under submerged culture, reverse transcribed into cDNA and *csgA* transcripts were detected by real-time PCR. cDNA copy number was determined by a standard curve generated from WT gDNA vs CT value and corresponded to 2.5 ng input RNA. Expression values were calculated as the ratio of log2 copy number relative to wild-type at 0 h. Each point is an independent biological replicate that is the average of 2 technical replicates. *: *p* ≤ 0.05 **(C)** Complementation assays between kanamycin-resistant wild type (PH1100), tetracycline-resistant wild type (PH2067), *csgA*, or *todK^++^* strains. Equivalent numbers of the indicated strains were mixed and developed on nutrient limited agar 5 days at 32 C. Cells were harvested, treated with heat and sonication to select for spores and equivalent proportions of each sample was spotted on rich media plates containing either oxytetracycline (red boxes) or kanamycin (blue boxes), respectively. Strains in red or blue text contained oxytetracycline- or kanamycin-resistance genes, respectively. Representative pictures are shown from two independent biological replicates each containing two technical replicates.

If lack of the extracellular C-signal was the only factor in preventing the *todK*^++^ strain from developing, we would expect that sporulation could be restored by co-development with wild-type cells (Kroos and Kaiser, 1987; Hoang et al., 2024). To first confirm that WT could rescue a *csgA* mutant, we mixed *csgA*::*Tn5* (oxytetracycline-resistant; tet^R^) cells with an equivalent number of either WT (kanamycin-resistant; kan^R^) or *csgA*::*Tn5* (tet^R^) cells and induced development on CF agar plates. After 120 h of development, heat- and sonication-resistant spores were harvested and then germinated on rich media containing either kanamycin or oxytetracycline. Co-development of *csgA* (tet^R^) and WT strains produced tet^R^ colonies, while no colonies were observed if *csgA* (tet^R^) was mixed with itself (Fig. 3C). To determine whether WT could rescue *todK*^++^ sporulation, the *todK*^++^ (kan^R^) strain was likewise induced to co-develop with an equivalent number of either WT (tet^R^) or *csgA* (tet^R^) cells. Importantly, co-development of WT (tet^R^) and *todK*^++^ (kan^R^) did not yield any kan^R^ colonies, suggesting WT could not rescue *todK*^++^ sporulation (Fig. 3C). As expected, co-development of *todK*^++^ (kan^R^) and *csgA* (tet^R^) strains produced neither kan^R^ nor tet^R^ colonies (Fig. 3C), because both strains lack CsgA (Fig 3A). This observation suggests that C-signal is not the only factor disrupted to prevent development in the *todK*^++^ strain.

### MrpC targets are downregulated by overexpressing TodK

C-signaling is thought to activate the FruA transcription factor to stimulate aggregation and sporulation during development (Ogawa *et al*., 1996; Ellehauge *et al*., 1998; Ueki and Inouye, 2003). We next examined FruA levels by immunoblot analysis. In WT cells, FruA accumulation increased ∼30-fold between 0 and 18 h after starvation, followed by an additional burst of accumulation at 30 and 36 h (Fig. 4A). In contrast, FruA accumulation in the *todK*^++^ mutant was severely reduced. FruA levels remained undetectable until 30 hours after which it accumulated to 60% of WT levels. In the *todK_H275A_*^++^ strain, FruA accumulation was similar to WT levels, consistent with the restoration of aggregation and sporulation. Furthermore, qRT-PCR analysis of RNA isolated from cells developing for 18 and 24 hours indicated a 14-fold reduction of *fruA* transcript in *todK*^++^ cells compared to WT (Fig. 4D). Thus, both C-signal and FruA are reduced if active TodK is overproduced.

**Fig. 4.**
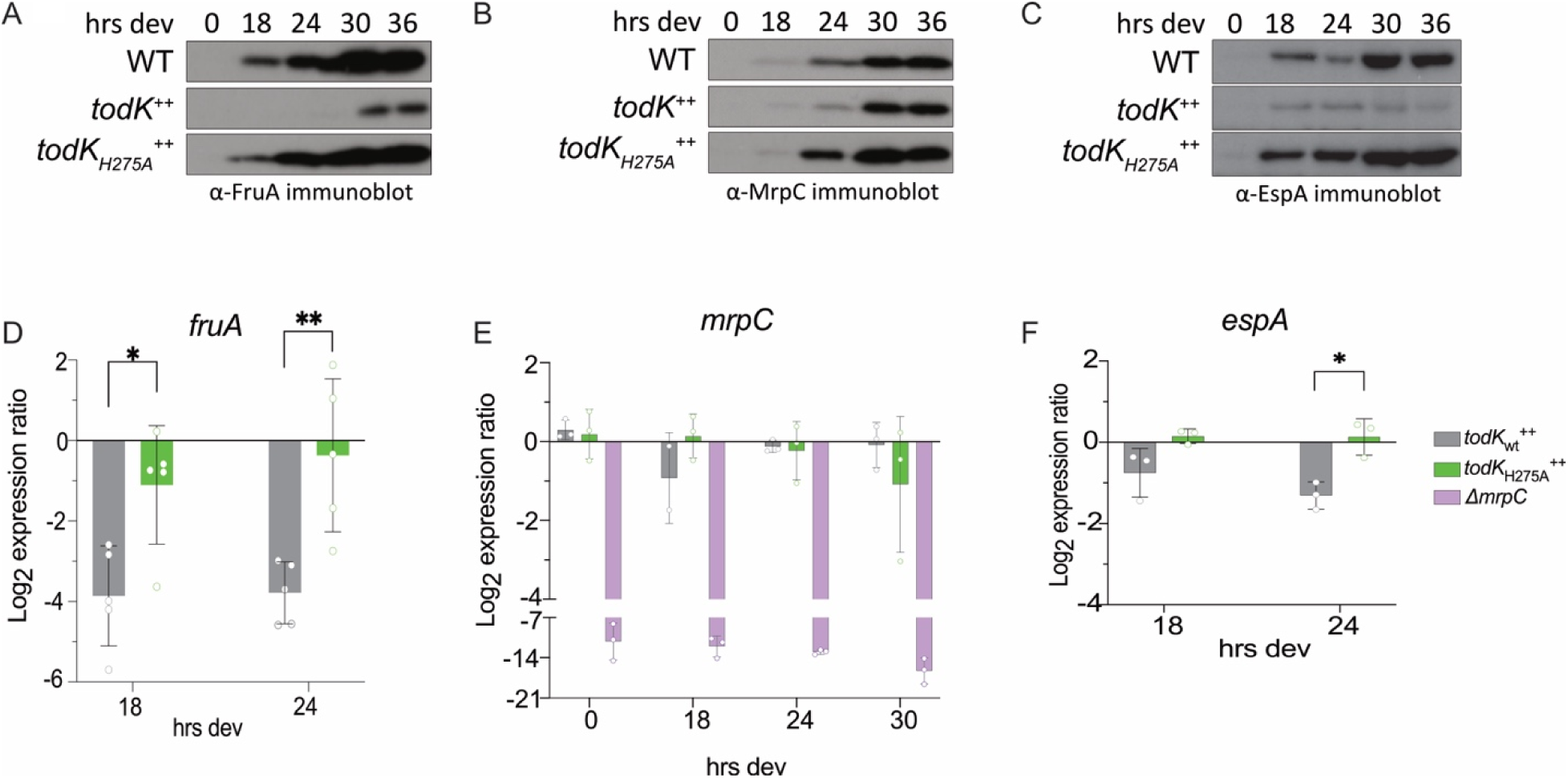
Targets of MrpC are downregulated in the *todK*^++^ strain. Immunoblot (**A-C**) or quantitative reverse transcriptase PCR (**D-F**) analysis of samples prepared from wild type (WT), *todK*^++^, and *todK_H275A_*^++^ strains developing under submerged culture. (**A-C**) Protein lysates (10 μg) were subject to immunoblotting with anti-FruA (**A**), anti-MrpC (**B**), or anti-EspA (**C**) antibodies. (**D-E**) RNA was isolated, transcribed to cDNA and detected with real-time PCR using primers to *fruA* (**D**), *mrpC* (**E**) or *espA* (**F**) Expression of each gene was calculated as using the ΔΔCT method using 16S CT values as an internal control (Livak and Schmittgen, 2001). Relative expression for each gene was calculated as the log2 ratio of the ΔΔCT values for *todK*^++^ or *todK* ^++^ Each point is an independent biological replicate that is the average of 2 technical replicates. **: *p* ≤ 0.01; *: *p* ≤ 0.05

Expression of *fruA* depends on the transcription factor, MrpC (Ueki and Inouye, 2003). We next examined MrpC accumulation by anti-MrpC immunoblot analysis (Fig. 4B). In WT cells, MrpC begins to rapidly accumulate by 24 h of development, which is necessary to stimulate aggregation (Sun and Shi, 2001b). Surprisingly, although the *todK*^++^ strain completely failed to develop, the MrpC accumulation pattern was not drastically different from WT. We observed a 2.7-fold reduction in MrpC accumulation at 24 hours of development, but by 30 and 36 h, MrpC was restored to WT levels. In the *todK_H275A_*^++^ strain, MrpC accumulation was 1.5-fold higher and 2.5-fold higher than WT at 18 h and 30 / 36h, respectively (Fig. 4B). qRT-PCR analysis did not reveal consistent significant differences in *mrpC* transcript levels between the WT, *todK*^++^, and the *todK_H275A_*^++^ strains (Fig. 4E). Together, these results strongly suggest that overexpression of active TodK somehow prevents MrpC from inducing appropriate *fruA* expression.

To determine if additional MrpC targets were affected by overexpression of TodK, we next examined several targets that are known to be dependent on MrpC, but not FruA. To confirm that CsgA fits this category, we examined accumulation of CsgA in developmental lysates (0-30h) prepared from *mrpC* or *fruA* mutants. As has been suggested previously, CsgA fails to accumulate in an *mrpC* mutant (Robinson *et al*., 2014; Farrugia *et al*., 2025), but displays WT levels in the *fruA* mutant (Saha *et al*., 2019b) (Fig. 3A). Second, we examined accumulation of the histidine kinase EspA which we have previously demonstrated is dependent on MrpC (Kasto and Higgs, 2025). Anti-EspA immunoblot analysis of developmental lysates from WT, *todK*^++^, and *todK_H275A_*^++^ strains indicated that EspA accumulation in the *todK*^++^ strain was significantly reduced compared to WT but was slightly increased in the *todK*_H275A_^++^ mutant (Fig. 4C). Additionally, qRT-PCR analysis confirmed that the *espA* transcript was also significantly decreased relative to WT in the *todK*^++^, but not the *todK_H275A_*^++^, strains (Fig. 4F). Thus, overexpression of catalytically active TodK prevents MrpC from activating multiple downstream targets.

## Discussion

TodK was previously characterized as a regulator of developmental timing because deletion of *todK* accelerates aggregation on nutrient-limited agar (Rasmussen and Sogaard-Andersen, 2003). Our newly developed rigorous and quantitative analysis of *todK* mutant development on NLA and SC revealed distinct phenotypes that provide insight into when and where TodK functions during development. A key phenotype shared between both conditions was the production of a greater number of fruiting bodies by the Δ*todK* mutant relative to WT (2.6-fold more on NLA and 3-fold more on SC) (Fig. 1D, Table 2, and Fig. 2D). This common effect suggests that TodK normally limits the number of initial aggregates that progress to fruiting bodies. One possible explanation is that TodK acts to delay the onset of sporulation (Fig. 1I) such that initial smaller aggregates can coalesce into larger structures before transitioning into mature fruiting bodies (Movie S1 vs S2). In contrast, TodK appears to have condition-dependent roles earlier in development. On NLA, TodK most strongly delays the transition to aggregation at the spot periphery, shows weaker activity in the spot core, and has little or no detectable effect on aggregation onset under SC conditions (Fig. 1E, S1). These spatial and environmental differences are consistent with the possibility that TodK is influenced by nutrient availability. Nutrient levels are likely highest at the periphery of the CF agar spot, lower in the core where dense cell populations consume available resources, and absent under the homogeneous starvation conditions of SC.

**Table 2.**
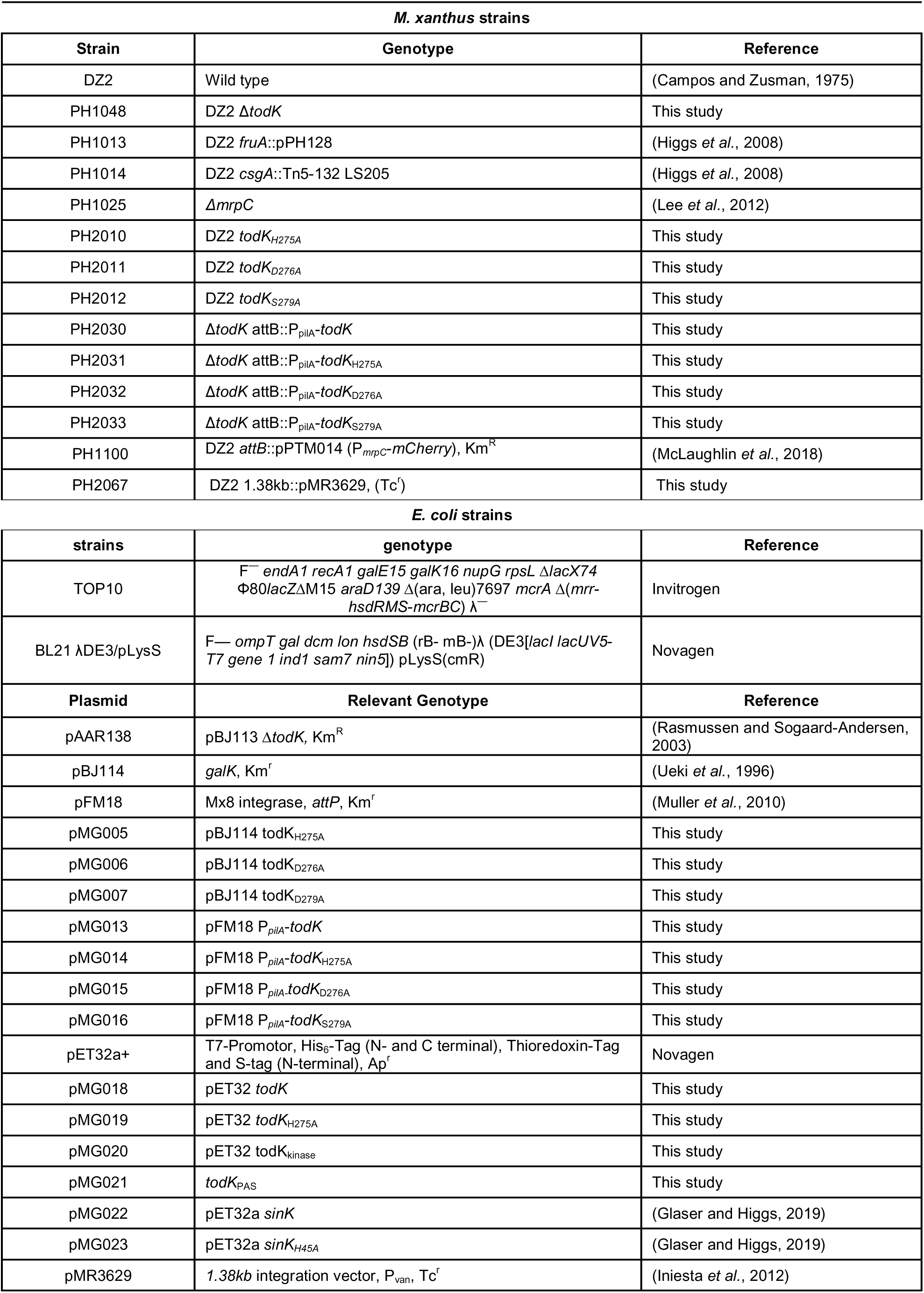
Strains and plasmids used in this work.

The striking complete inhibition of aggregation observed by overexpression of wild type TodK (Fig. 2A) is consistent with the reduced inhibition of developmental stages in the Δ*todK* mutant. These findings further suggest that TodK abundance itself is a major determinant of developmental inhibition and may, when sufficiently elevated, override normal regulatory inputs. This interpretation is consistent with the apparently low abundance of chromosomally encoded TodK. TodK is barely detectable in wild type whole-cell lysates (Fig. 2B) despite the high sensitivity of our TodK antisera relative to anti-sera we generated against other HKs (Schramm, 2012; Schramm *et al*., 2012; Glaser, 2017), suggesting that the protein is either present at very low levels throughout the population or restricted to a subpopulation of cells. In either case, TodK is likely normally present at substantially lower stoichiometry than its target(s), consistent with both the modest phenotype of the Δ*todK* mutant and the dramatic developmental arrest caused by forced expression throughout the population.

Beyond revealing the importance of TodK dosage, the overexpression strain also provided insight into the downstream pathway through which TodK regulates developmental transitions. Most notably, *mrpC* expression remained largely unchanged in the *todK*^++^strain (Fig. 4B and E), indicating that the starvation-sensing and population-dependent signaling pathways required for *mrpC* induction remained functional. In contrast, expression of several MrpC-dependent targets, including *csgA*, *fruA*, and *espA*, was strongly reduced (Figs. 3 and 4). MrpC is an attractive target to control developmental transitions, because its activity is necessary to drive the distinct thresholds of C-signal-dependent FruA accumulation initially proposed to be the TodK target (Rasmussen and Sogaard-Andersen, 2003), and which stimulate first aggregation and then sporulation (Lobedanz and Sogaard-Andersen, 2003; Hoang *et al*., 2021).

Collectively, our observations support a model in which TodK regulates distinct stages of developmental progression by constraining the activity of MrpC. The environment-dependent effect of TodK on aggregation timing suggests that its role during early development is modulated by nutritional conditions. In contrast, the accelerated onset of sporulation resulting in the production of a larger number of smaller fruiting bodies observed in the Δ*todK* mutant under both NLA and SC conditions indicates a second, more broadly conserved role in regulating progression from aggregation to sporulation. Thus, TodK appears to regulate two distinct developmental checkpoints: a nutrient-sensitive checkpoint controlling aggregation onset and a second checkpoint controlling entry into sporulation.

The molecular targets that link TodK to MrpC inhibition remain unknown. Because purified MrpC binds target promoters readily *in vitro* and no activating ligand has been identified, TodK likely influences MrpC through post-translational regulation. MrpC has previously been proposed to be regulated by Ser/Thr kinase pathways involving Pkn14 (Nariya and Inouye, 2006; Feeley *et al*., 2019). However, development is not restored in a *todK*^++^ *pkn14*_K48N_ double mutant (data not shown), arguing that TodK does not act solely through the Pkn14 pathway. Alternatively, TodK may regulate proteins that interact directly with MrpC. The hybrid histidine kinases AskA and AskB are attractive candidates because they physically associate with MrpC and are required for normal developmental regulation (Marcos-Torres *et al*., 2020). Although AskA and AskB contribute to proper induction of *mrpC* expression, which is largely unaffected in the *todK*^++^ strain, the functional consequences of their physical interactions with MrpC remain incompletely understood. Regardless, we favor a model in which TodK post-translationally regulates the activity of accumulated MrpC rather than its transcriptional induction. Identification of the immediate TodK target(s) will be required to clarify this mechanism.

The severe TodK-overexpression phenotype also enabled genetic analysis of TodK signaling states. Unexpectedly, both mutations predicted to render TodK kinase-defective phosphatase-active and kinase-active phosphatase-defective restored development when overexpressed (Fig. 2). Similarly, equivalent mutations introduced at the native chromosomal locus largely phenocopied the Δ*todK* mutant (Fig. S2). Under a conventional two-component signaling model, disruption of either kinase or phosphatase activity would be expected to bias signaling toward one regulatory state rather than abolish TodK-dependent inhibition. These findings are inconsistent with a simple model in which TodK functions exclusively as either a kinase or phosphatase on a single target. Instead, they suggest that both catalytic activities are required for proper signaling. One possibility is that TodK phosphorylates one substrate while dephosphorylating a second substrate that exerts an opposing effect on MrpC activity. In such a model, disruption of either catalytic branch would prevent TodK-dependent inhibition of MrpC. Although speculative, this hypothesis is consistent with precedents in which a histidine kinase acts on distinct response regulators through different catalytic activities. For example, in the *E. coli* nitrate response, the HK NarX preferentially acts as a kinase on the RR NarP, but primarily acts as phosphatase on the RR NarL∼P (Rabin and Stewart, 1993; Williams and Stewart, 1997).

Finally, the observation that increased TodK activity can completely abolish formation of the *M. xanthus* multicellular biofilm highlights the remarkable capacity of histidine kinases to control global regulatory networks and developmental fate decisions. Histidine kinases are attractive antimicrobial targets because they regulate numerous pathogenic behaviors yet are absent from higher eukaryotes (Chen *et al*., 2022). The robust TodK-overexpression phenotype may provide a tractable experimental platform for identifying small molecules that modulate histidine kinase activity and for defining the environmental signals that regulate TodK-dependent developmental control.

## Materials and Methods

### Bacterial strain and growth conditions

The bacterial strains in this study are listed in Table 2. *Escherichia coli* strains were grown under standard laboratory conditions in LB medium supplemented with 50 μg ml^−1^ kanamycin or 20 μg ml^−1^ tetracycline where necessary (Sambrook et al., 1989). *M. xanthus* strains were grown at 32 °C in Casitone Yeast Extract (CYE) broth (Campos and Zusman, 1975) with shaking at 220 rpm, or on CYE-1.5% agar plates supplemented with kanamycin at 100 μg ml^−1^ or oxytetracycline at 10 μg ml^−1^, where necessary.

### Construction of plasmids and strains

Strains used in this study and the plasmids used to generate the deletion of or point mutations in the *todK* genomic locus are listed in Table 2. Plasmids were constructed as described in detail previously (Lee *et al*., 2010) using the primers listed in Table S1. Strains PH1048, PH2010, PH2011, and PH2012 were constructed by introducing plasmids pAAR138, pMG005, pMG006, or pMG007, respectively, into *M. xanthus* strain DZ2 by electroporation followed by selection (kanamycin) and then counter-selection (galactose) and then PCR-based confirmation of the respective mutation using the primers listed in Table S1, as described in detail previously (Lee *et al*., 2010). Plasmids for overexpression of *todK* genes were constructed using overlap PCR to fuse *todK* alleles, amplified from genomic DNA of the wild type or respective *todK* point mutations, to the *pilA* promoter using the primers and vectors indicated in Table S1.

Overexpression strains PH2030, PH2031, PH2032, or PH2033 were generated by introduction of plasmids pMG013, pMG014, pMG015, or pMG016 into the PH1048 strain and site-specific integration into the genomic Mx8 *attB* site (Magrini *et al*., 1999) was selected by resistance to kanamycin and confirmed by PCR screen using primers described in Table S1, as described previously (Lee *et al*., 2010; Feeley *et al*., 2019).

### Developmental Assays

To induce development on nutrient-limited agar plates, 1 ml overnight cultures of *M. xanthus* were harvested, washed in MMC buffer (10 mM MOPS pH 7.6, 4 mM MgSO_4_, 2 mM CaCl_2_), resuspended in MMC to A_550_ 7 (∼ 4 x10^9^ cells ml^-1^), and 10 μl were spotted on Clone-Fruiting (CF) agar plates (10 mM MOPS pH 7.6, 8 mM MgSO_4_, 1 mM KH_2_PO_4_, 0.2 % sodium citrate, 0.02% ammonium sulphate, 1.5% agar, 0.1% sodium pyruvate). After drying, plates were inverted and incubated at 32 °C for 72-120 hours. Pictures were recorded using a Leica MZ8 stereomicroscope and Leica DFC320 camera.

Development of *M. xanthus* strains in submerged culture (SC) conditions was performed as described in detail previously (Lee et al., 2010). Briefly, vegetative cells from broth cultures were diluted to an optical density of 0.035 at 550 nm (∼ 2 x 10^7^ cells ml^−1^) in CYE broth and 0.5, 0.15 or 16 ml were added to 24-well tissue culture, 96-well tissue culture, or 100 mm petri dishes and incubated for 24 hours at 32°C. To induce development, CYE was replaced with an equivalent volume of MMC and incubated at 32°C for the indicated times.

Video analysis of development was performed in submerged culture on 96-well plates as described previously (Glaser and Higgs, 2019). Briefly, the center of each well was imaged every 30 min for 72 h in a Tecan Spark 10M plate reader and the resulting images were compiled into movies at 6 frames per second. For each movie generated, the first frame where visible aggregates were observed was recorded as aggregation onset. Maximum aggregates were totaled from the frame wherein the most aggregation centers were observed. Fruiting bodies were enumerated from the final frame of each video. Data were compiled from three independent biological experiments, each with 3 replicates per strain. Statistical significance was determined for relevant comparisons by Welch’s t-test.

### Sporulation Assays

To enumerate total numbers of heat- and sonication-resistant spores produced, *M. xanthus* strains developed under submerged culture in 24-well tissue culture plates. For each time point, cells were harvested from triplicate wells, pelleted for 5 min at 17 000 × g, the supernatant was removed, and the pellets were stored at −20°C until further use. To determine sporulation efficiency, pellets were resuspended in 0.5 ml of sterile water, heated at 50°C for 60 min, and sonicated at 30% output for 15 s (0.5-s pulses) using a microtip and Branson Sonifier 250 (Branson Ultrasonics). Phase-bright spores were enumerated with a Helber bacteria counting chamber (Hawksley, UK). Average total spores and associated standard deviations were reported from three independent biological replicates. Statistical significance was determined by unpaired two-tailed t-tests.

### Image Analysis of Nutrient Limited Agar (NLA) developmental phenotypes

The image-analysis pipelines were implemented in Python with code-generation assistance from a large language model (Claude; Anthropic, San Francisco, CA). All analytical logic, parameters and outputs were designed, verified and validated by the authors against the source images; no generative AI was used to produce or modify the underlying image data or the reported measurements. Detailed steps for each pipeline are provided in Supplemental Methods. All parameter values are itemized in the accompanying parameter provenance table (Supplemental Methods, Table S2). The image-analysis code used in this study is archived at Zenodo (DOI: 10.5281/zenodo.21538883) and on GitHub (https://github.com/stuarthuntley/Myxococcus-cf-image-analysis), released under the PolyForm Noncommercial License 1.0.0, which permits non-commercial academic use and reproduction. Commercial rights are reserved by the authors.

Images of developmental spots (2048 × 1536 px RGB brightfield stereomicrographs) were first converted to grayscale and a spatial scale was set as 0.005123 mm/px (5.123 µm/px) at full resolution. Analyses were implemented in Python 3.11.4. Image processing used OpenCV 4.13.0.92 (Bradski, 2000) for the texture pipeline and scikit-image 0.26.0 (Van Der Walt *et al*., 2014) for blob detection and morphology; numerical work used NumPy 2.4.6 (Harris *et al*., 2020); statistical tests used SciPy 1.17.1 (Virtanen *et al*., 2020). Radial density figures were produced with Matplotlib 3.11.0 (Hunter, 2007). The interactive circle-setting step was performed in a purpose-built browser-based editor. The editor was a local HTML/JavaScript tool in which the operator overlaid an adjustable circle on each spot image to set the spot center and growth-front radius; accepted values were exported verbatim for downstream analysis, with no automatic refit.

### Aggregation analysis

For aggregation analysis, a texture (granularity) metric proxy was developed. Granularity is defined as the standard deviation of a band-pass (difference-of-Gaussians) filtered, flat-field-corrected image within a fixed circular region of interest. The grayscaled image was first downscaled by two using an averaging resize. Illumination was corrected by dividing by a heavily blurred copy of itself (σ = 60 half-resolution px) rescaled to the background mean. A difference-of-Gaussians band-pass (σ = 1 minus σ = 6) isolated fruiting-body-scale structures; and the population standard deviation of that band-passed image was taken within the ROI. Because the operation was a band-pass, the metric was insensitive both to the smooth illumination gradient and to absolute image brightness, responding only to structure at the scale of interest. The ROI was a fixed 720 px (full-resolution) radius circle — identical for every spot, since the spots were equal-volume drops of similar size — centered on an automatic drop-edge circle fit. Three regions were reported: whole spot (0–1.0 R), inner core (0–0.65 R), and outer ring (0.65–1.0 R, the spot periphery). Values were raw standard deviations in arbitrary units with no scaling applied. Normalization to the same-experiment wild-type baseline was a separate downstream step applied to per-strain means, not baked into the stored values. Statistically significant differences were assessed by paired t-test on per experiment means (n= 6) for DZ2 vs Δ*todK* or n =3 for the *todK* point mutants.

### Spatial organization

For fruiting body spatial organization analysis, a profile of darkness against radius was developed. Darkness was measured as local-background-referenced intensity: a Gaussian estimate of the local background (σ ≈ 100 full-resolution px, ≈ 0.5 mm) was subtracted from the grayscale image and the result clipped at zero. A local reference ensured that spatially broad features (i.e. bare agar, the illumination gradient, and the underlying cell lawn) were absorbed into the background and removed, so that only structures compact and dark relative to their immediate surroundings survived. The metric therefore reports each body’s darkness in excess of the local cell and agar background.

For each spot, the center and the growth-front radius were set by hand using the interactive circle editor (above), and used verbatim with no automatic refit. The profile was the azimuthal (360-degree) mean darkness in 1-px annuli measured outward from the operator-defined spot center, smoothed with a 45-px moving average, and plotted against absolute radial distance from the center (mm; 0.005123 mm/px). The operator-set growth-front radius was used only to mark the colony edge and to define the coverage cutoff (below), not to rescale the axis. The fraction of each annulus lying inside the image frame (coverage) was tracked; profiles were drawn solid while coverage is complete and dotted once the sampling ring ran off the frame.Profiles represent the mean of six independent biological replicates; shaded band, SEM. Profiles are presented descriptively; no hypothesis test was applied to the curves.

Spatial analysis of fruiting bodies was performed by Laplacian-of-Gaussian blob detection on the local-background darkness map (not on the raw image), with a darkness gate that rejected candidates failing an absolute darkness floor, followed by size-ordered nesting resolution and a 1/6-overlap merge rule that fused candidate detections overlapping by more than one sixth. Detections were eye-validated against the images. The detector emitted, per body, a center (x, y) and a fitted circle radius.

Spatial statistics were then computed per spot: count; density (bodies per mm² of spot area); and the mean nearest-neighbor distance (NND). Statistically significant differences were assessed by a two-sided Mann–Whitney U on the six per-experiment means (6 versus 6), because the two strains had grossly unequal spread.

### Overexpression and purification of recombinant protein

pET32a plasmids (Novagen) carrying *todK*_WT_ (pMG018) or *todK*_H275A_ (pMG019) were induced in *E. coli* BL21lDE3/pLysS cells as follows: Cultures were grown with shaking (220 rpm) to mid-logarithmic phase (OD_600_ ∼0.5) at 37 °C and transferred to 18 °C until OD_600_ ∼0.7. Expression was induced with 0.5 mM IPTG followed by incubation overnight at 18 °C. Cultures were pelleted and stored at -20 °C. Recombinant proteins were purified as described previously in detail (Glaser and Higgs, 2019). Briefly, pellets were resuspended in binding buffer, disrupted by French press, and clarified (3,500 × g for 15 min 4°C then 20,000 × g for 30 min at 4 °C). Trx-His_6_-TodK and Trx-His_6_-TodK_H275A_ proteins were purified in a fast protein liquid chromatography (FPLC) affinity system (Äkta Technology, GE Healthcare) using a 1 ml HisFF1 trap nickel affinity column (GE Healthcare) with a flow rate of 1 ml/min at 4 °C. The column was washed of unbound protein, and TodK proteins were eluted a gradient of imidazole (20 nM to 500 nM). Fractions were analyzed by denaturing polyacrylamide gel electrophoresis (SDS-PAGE) for purity and concentration. Fractions containing peak Trx-His_6_-TodK and Trx-His_6_-TodK_H275A_ proteins were pooled and dialyzed overnight at 4 °C against storage buffer. Trx-His_6_-SinK proteins were purified as described in detail previously (Glaser and Higgs, 2019).

### Generation of anti-TodK immunosera

Rabbit anti-TodK antibodies were generated by Eurogentec (Belgium) using the 28-day immunization program. Anti-TodK antibodies were affinity purified from immune serum using purified Trx-His_6_-TodK as described previously (Glaser and Higgs, 2019).

### Cell lysate preparation and immunoblot analysis

For immunoblot analysis of developing cell lysates, *M. xanthus* strains were induced to develop under submerged culture conditions in 100-mm petri dishes (Lee *et al*., 2010). At the indicated time points, cells were harvested and proteins were precipitated by addition of trichloroacetic acid (TCA), solubilized in protein sample buffer, and adjusted to 1 μg μl^−1^ in sample buffer as described in detail previously (Schramm *et al*., 2012). 10 μg of each sample was resolved by denaturing 11% (CsgA, MrpC, and FruA) or 8% (TodK and EspA) PAGE and transferred to polyvinylidene fluoride (PVDF) membranes (Millipore Sigma) using a semidry transfer apparatus (Hoefer). Immunoblot analyses were performed using anti-TodK (this study), anti-CsgA, anti-FruA (Lobedanz and Sogaard-Andersen, 2003), anti-MrpC (Lee *et al*., 2012), or anti-EspA (Higgs *et al*., 2008) antibodies at 1:1000 (TodK, MrpC, EspA) or 1:500 (CsgA, FruA) dilutions. Goat anti-rabbit IgG secondary antibodies conjugated to horseradish peroxidase (HRP; Pierce) were used at a dilution of 1:20 000, and signal was detected with enhanced chemiluminescence substrate (Thermo Fisher Scientific), followed by exposure by iBright™ FL1500 Imaging System (Invitrogen) (TodK) or exposure by autoradiography film (Kodak) (CsgA, FruA, MrpC, and EspA).

### In vitro phosphorylation assays

10 μM recombinant affinity tagged TodK, TodK_H275A_, SinK, or SinK_H45A_ were added to phosphorylation reaction buffer (storage buffer supplemented with 50 mM KCl and 5 mM MgCl_2_). Where indicated, MgCl_2_ was increased to 10 mM or substituted with 1 or 10 mM MnCl_2_ or CaCl_2_. Autophosphorylation reactions were initiated by addition of 0.5 mM ATP and 1.7 μM [γ-^32^P]-ATP (222 TBq/mmol, Hartmann Analytic, Braunschweig) and the reaction was incubated for 0 – 60 min at 32 °C. For each time point, a 10 μl sample was removed from the reaction mix and quenched with 20 μl 2 x LSB. Samples were immediately stored at -20 °C. Samples were resolved on a 12 % polyacrylamide gel by SDS-PAGE, exposed to a phosphorimager screen (GE Healthcare) over night, and analyzed using a StormTM 800 imaging system (GE Healthcare). To confirm equal loading, gels were subsequently stained using Coomassie blue.

### Quantitative real-time PCR

mRNA levels were examined using quantitative real-time PCR. *M. xanthus* strains were induced to develop under submerged culture conditions in a 100-mm petri dishes. The cell layer was resuspended in 0.5 ml MMC and 1.5 ml RNAprotect Bacteria Reagent (Qiagen) and incubated at room temperature (RT) for 5 min. Samples were pelleted at 17000 × g for 1 min, flash frozen in liquid nitrogen, and stored at −80°C until further use. Total RNA was isolated using carbon-coated magnetic beads that isolate RNA from gDNA using a CarbonPrep Tissue Kit (Life Magnetics, LM) per manufacturer’s instructions. Briefly, pellets were thawed and resuspended in lysis solution (4:1 lysis buffer stock and DEPC-treated water, lysozyme [10 mg ml^−1^], and proteinase K [10 mg ml^−1^]). RNA Pre-Binding solution (RPS) was added to the lysates followed by LM magnetic beads. Samples were placed on magnetic stands and supernatant was removed from beads. The beads were washed with RPS-Wash buffer and then twice with Wash Buffer and allowed to dry on magnetic stands. RNA was recovered from the beads in DEPC-treated water by incubation at 65°C for 5 min to disrupt RNA-bead complex. Samples were pelleted (10000 × g for 1 min) and supernatant containing total RNA was stored at −80°C until further use.

1-2 μg RNA sample was treated with DNase, and 500 ng RNA was transcribed into cDNA using random hexamer primers as described in detail previously (Higgs *et al*., 2008). RT-PCR was performed in a QuantStudio 3 Real-Time PCR System (Applied Biosystems) using PowerUp SYBR green PCR master mix (Applied Biosystems) and *fruA* (oPH252/oPH253), *mrpC* (oPH353/oPH354), *espA* (oPH369/oPH370), or 16S rRNA (oPH235/oPH236) primers. No template controls contained DEPC-treated water instead of cDNA template. Standard curves generated from 10, 1, 0.1, and 0.01 ng of wild-type genomic DNA (gDNA) were analyzed to calculate primer pair efficiency, and for calculating approximate gene copy number. Real-time PCR cycles were performed as described previously (Kasto and Higgs, 2025). Threshold cycle (C_T_) values for each reaction were assigned automatically by the system software (QuantStudio Design & Analysis v1.5.1). Average CT values were calculated from two technical replicates.

Standard curves were generated by converting ng of gDNA to log_10_ transformed DNA copy number and plotting experimental gDNA C_T_ values against the respective log_10_ transformed DNA copy number.A linear regression for each standard curve was generated, and each experimental gene C_T_ value was fitted to the respective linear regression to calculate approximate gene copy number per reaction. Relative gene expression levels were calculated using the comparative C_T_ method using 16S C_T_ values as the internal control. Expression values are presented as log_2_ ratio of either the DNA copy number (*csgA*) or the gene expression level *(fruA*, *mrpC*, *espA*), relative to the WT DZ2 sample at t=0 post starvation.

### Complementation assays

Overnight vegetative cultures were washed and resuspended to 4 x 10^9^ cells mL^-1^ in MMC buffer.Concentrated cells were mixed pairwise at a 1:1 ratio and triplicate 20 µL (containing 4 x 10^8^ cells of each strain) aliquots were spotted on CF agar and incubated at 32 °C for 5 days.For each sample, cells were scraped of the agar surface, resuspended in 0.5 ml ddH_2_O, and treated by heat and sonication as described above. 10 ul spots were plated on CYE oxytetracycline or CYE kanamycin plates, followed by followed by incubation at 32 °C for at least 5 days. *csgA* or *todK++* strains were scored as complemented if spores produced germinated into colonies on oxytetracycline or kanamycin containing plates, respectively. Strains were scored as failed to complement if no growth was observed on the respective plates. Complementation results were consistent across three independent biological replicates.

## Acknowledgments

This work was supported by funds from the National Science Foundation grant IOS-1651921 (PIH), the Max Planck Society (PIH), and a fellowship from the International Max Planck Research School (MG). We thank Dr. Bongsoo Lee for the construction of PH1048 (DZ2 Δ*todK*), the Greenberg lab (Wayne State University) for use of the iBright imaging system, and Drs. Shelby Kasto, Maeve McLaughlin, and Brooke Feeley for helpful discussions. We additionally acknowledge the assistance of the Wayne State University Proteomics Core that is supported through NIH grants P30ES036084, P30CA022453 and S10OD030484.

