## Supplemental data for "The orphan histidine kinase TodK controls *Myxococcus xanthus* biofilm development by inactivating the CRP/Fnr homolog, MrpC"

**Movies S1.** Developmental phenotype of the wild-type (WT) strain, DZ2.

**Movies S2.** Developmental phenotype of the Δ*todK* strain

**Movies S3.** Developmental phenotype of the *todK*^++^ strain

**Movies S4.** Developmental phenotype of the *todK_H275A_*^++^ strain.

**Movies S5.** Developmental phenotype of the *todK_D276A_*^++^ strain.

**Movies S6.** Developmental phenotype of the *todK_S279A_*^++^ strain

**Fig. S1** Image analysis tools for quantification of developmental phenotypes on nutrient-limited agar.

**Fig. S2.** Predicted kinase and/or phosphatase mutations generated in the native *todK* locus display ∆*todK*-like CF developmental phenotypes.

**Fig. S3.** TodK autophosphorylation cannot be detected *in vitro*.

**Table S1.** Primers used in this work

**Supplemental Methods:**

**Table S2.** NLA image-analysis pipeline parameters

**Detailed NLA image analysis methods**

Movie S1. Developmental phenotype of the wild-type (WT) strain, DZ2. A representative movie of WT DZ2 *M*. *xanthus* cells imaged by a high-resolution submerged culture assay. Cells were induced to develop in 96 well plates and images were recorded every 30 min from 0 to 72 hours post starvation and compiled using ImageJ.

Movie S2. Developmental phenotype of the Δ*todK* strain. A representative movie of strain PH1048 imaged by a high-resolution submerged culture assay. Cells were induced to develop in 96 well plates and images were recorded every 30 min from 0 to 72 hours post starvation and compiled using ImageJ.

Movie S3. Developmental phenotype of the *todK*^++^ strain. A representative movie of strain PH2030 imaged by a high-resolution submerged culture assay. Cells were induced to develop in 96 well plates and images were recorded every 30 min from 0 to 72 hours post starvation and compiled using ImageJ.

Movie S4. Developmental phenotype of the *todK_H275A_*^++^ strain. A representative movie of strain PH2031 imaged by a high-resolution submerged culture assay. Cells were induced to develop in 96 well plates and images were recorded every 30 min from 0 to 72 hours post starvation and compiled using ImageJ.

Movie S5. Developmental phenotype of the *todK_D276A_*^++^ strain. A representative movie of strain PH2032 imaged by a high-resolution submerged culture assay. Cells were induced to develop in 96 well plates and images were recorded every 30 min from 0 to 72 hours post starvation and compiled using ImageJ.

Movie S6. Developmental phenotype of the *todK_S279A_*^++^ strain. A representative movie of strain PH2033 imaged by a high-resolution submerged culture assay. Cells were induced to develop in 96 well plates and images were recorded every 30 min from 0 to 72 hours post starvation and compiled using ImageJ.

Fig S1

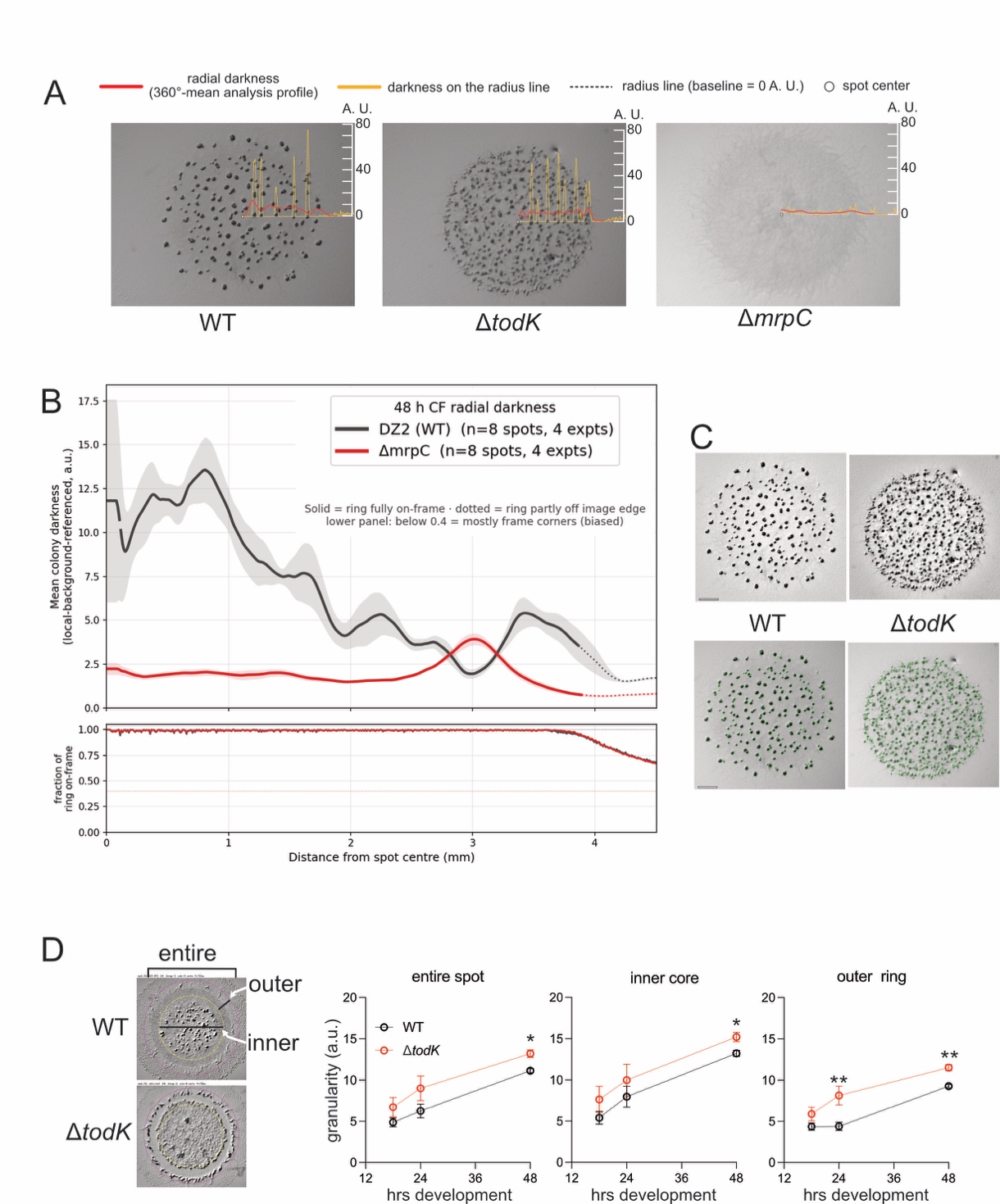

**Fig. S1. Image analysis tools for quantification of developmental phenotypes on nutrient-limited agar.** (**A**) Depiction of radial density analysis to detect organized fruiting body patterns in wild type (WT; DZ2), Δ*todK* (PH1048), and the non-developing Δ*mrpC* (PH1025) strains developed on CF agar for 48h. A radial line (dotted) was generated from spot center (white circle), and background subtracted pixel density in arbitrary units (a.u.) is measured. For illustrative purposes, single-width pixel intensity measured along the line reflects fruiting bodies present. The 360° average pixel density is demonstrated by the red line. To detect consistent patterning, mean densities from multiple images were plotted against radial length in mm (Panel B). (B) Radial density analysis of the wild type (black lines) vs Δ*mrpC* (red lines) strains. Upper panel: Mean radial densities from 8 images collected from four independent biological experiments. Solid line, full 360-degree coverage; Dotted line, partial coverage; Shaded band, standard error of the mean. The non-developing mutant demonstrates little density accumulation throughout with a slight peak at the spot periphery (3 mm) which represents the colony front. Bottom panel: Portion of the circular spot within the rectangular image frame. Density analysis greater than 3.6 mm from the spot center reflects incomplete coverage at the rectangular image edges. (**C**) Analysis of fruiting body enumeration. Top images: Representative contrast-enhanced images of WT or Δ*todK* strains developed for 48 h spots on CF agar. Bottom images: The same two images with detected fruiting bodies circled (green), illustrating the automated detection used for the spatial statistics. Scale bar 1 mm. (**D**) Quantification of aggregation by image texture analysis. Left images: Representative contrast-enhanced images of WT or Δ*todK* strains developed for 24 h on CF agar. Texture (granularity) was measured for the entire spot (pink ring), for the inner core (yellow ring) or for the outer spot periphery (between pink and yellow rings) as indicated on the images. Right panels: Granularity detected at 18-, 24-, and 48-hour images for the three regions in the WT (black) or Δ*todK* (red) strains. The outer ring (spot periphery) panel is the same as Fig. 1E. The Δ*todK* mutant displays a trend for higher granularity compared to the wild type at all time points and in all regions, but it is significantly advanced in the spot periphery (outer ring) compared to wild type starting at 24 hrs development.

Fig S2

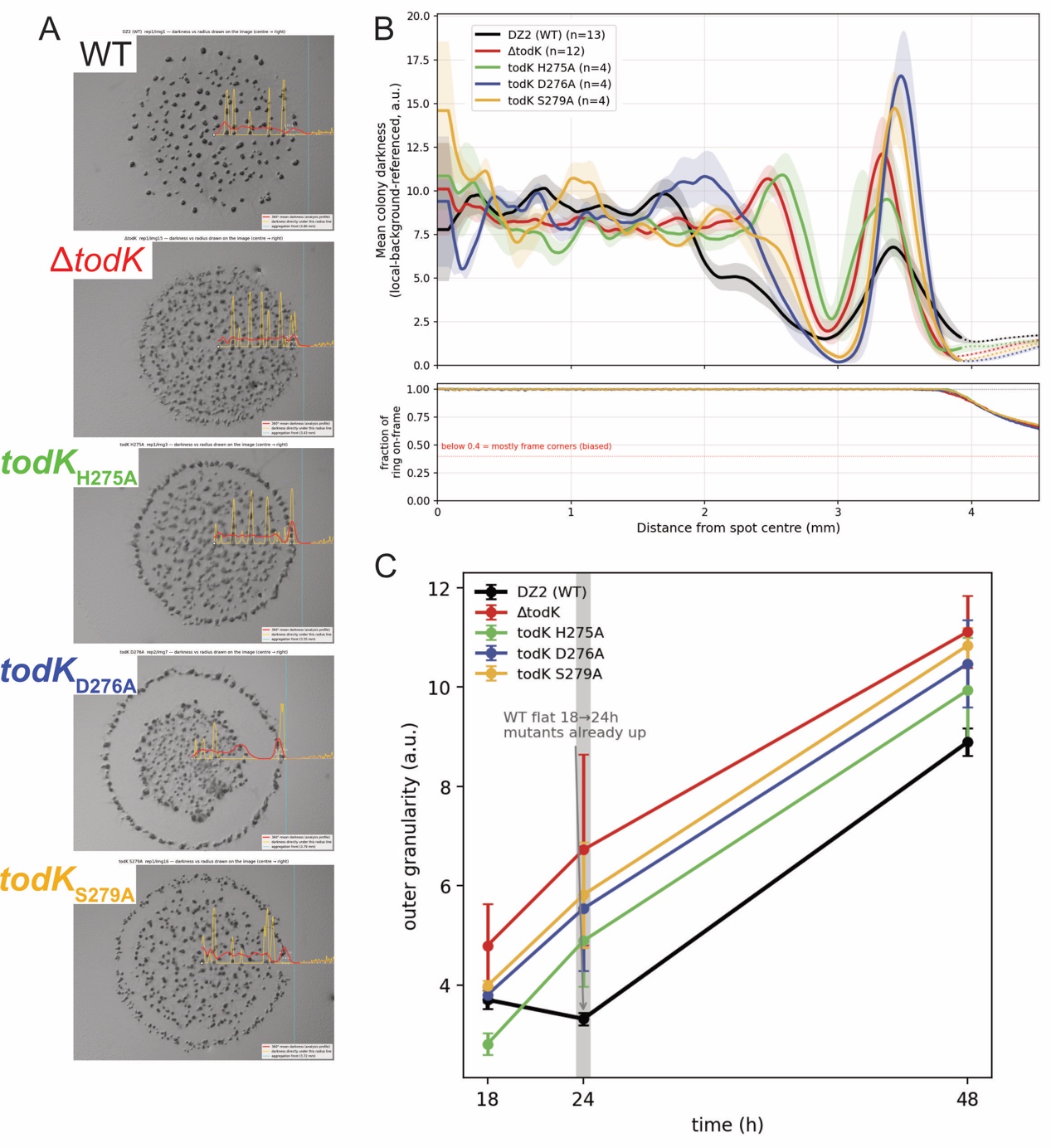

**Fig. S2. Predicted kinase and/or phosphatase mutations generated in the native *todK* locus display ∆*todK*-like CF developmental phenotypes.** (**A**) Representative fruiting body patterns produced by WT (DZ2), ∆*todK* (PH1049), *todK_H275A_*; K^-^ P^-^ (PH2010), *todK_D276A_*; K^-^P^+^ (PH2011), or *todK_S279A_*; K^+^P^-^ (PH2012) strains at 48 hours of development. Single- and mean 360°- pixel intensities for the respective images are depicted by yellow and red lines respectively. See Fig. S1A figure legend for details. (**B**) Average 360°- pixel intensities for the indicated mutants from at least 4 independent biological replicates. See Fig. S1B figure legend for details. (C) Spot periphery aggregation. The indicated *todK* point mutations, ∆*todK*, and the wild type strain were developed on CF agar and outer ring texture (granularity) was detected on 18-, 24-, and 48-hour images. All *todK* mutants display increased granularity at 24 hours compared to the wild type. Note the putative K+P- and K-P+ mutant alleles display larger body-free zones between the spot periphery and inner core.

Fig S3

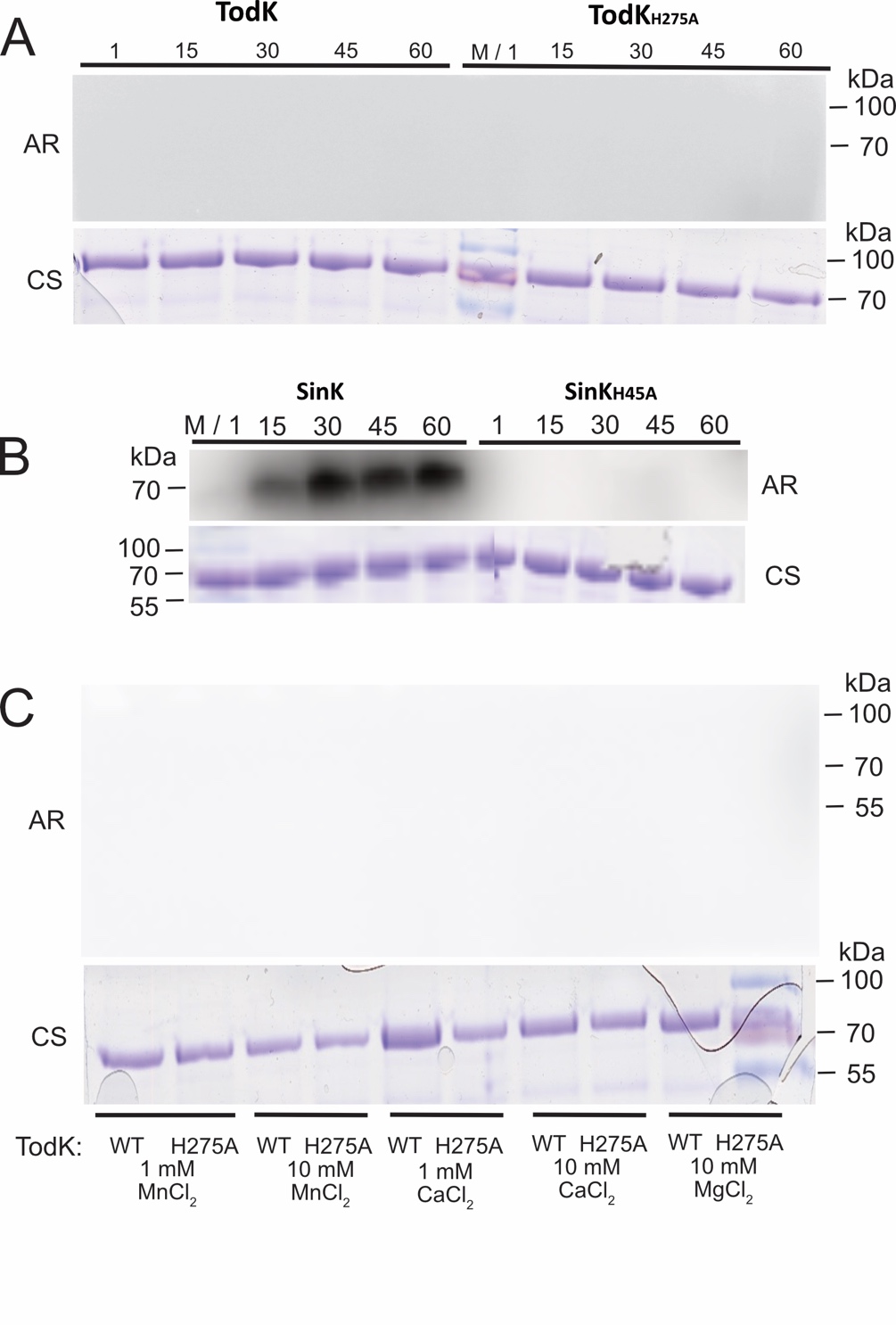

**Fig. S3. TodK autophosphorylation cannot be detected *in vitro*.** (**A**) TodK autophosphorylation assay. Autoradiograph (AR) and corresponding Coomassie blue-stained gel (CS) of 10 μM Trx-His_6_-TodK or Trx-His_6_-TodK_H275A_ incubated in the presence of [γ-^32^P]-ATP for the indicated minutes, resolved by SDS-PAGE, and exposed to a phosphoimager screen. (**B**) SinK autophosphorylation assay performed in parallel with A. Autoradiograph and corresponding Coomassie blue-stained gel of 10 μM Trx-His_6_-SinK or Trx-His_6_-SinK_H45A_ incubated in the presence of [γ-^32^P]ATP and detected as in A. (**C**). TodK autophosphorylation assay performed as in A with the indicated alternate cations replacing magnesium.

| **Table S1.** Primers used in this work | | | | | | | |
| --- | --- | --- | --- | --- | --- | --- | --- |
| Plasmid or Strain Name | Description^a^ | Oligo name | Primer type^b^ | Sequence (5'-3') | Restriction Site | Template DNA | Target Plasmid |
| pMG005 | pBJ114 todK_H275A_ | oPH972 | A | CG**GAATTC**CTGGTGCACTGCCGTGAC | EcoRI | DZ2 | pBJ114 |
|  |  | oPH973 | B | GTCCGCGGTGACGATGCCCATGAGCTG |  |  |  |
|  |  | oPH974 | C | ACCGCGGACATCCGCAGCCCGCTGGGC |  |  |  |
|  |  | oPH975 | D | CA**GGATCC**GAAGGGCTCGAAGAGGTG | BamHI |  |  |
| PH2010 |  | oPH992 | E_wt_ | GCTCATGGGCATCGTCACCC |  | todK_H275A_ loop-out clones | N/A |
|  |  | oPH993 | E_mut_ | GCTCATGGGCATCGTCACCG |  |  |  |
|  |  | oPH994 | F | CACCGTCTTGTTGCTCCAGCG |  |  |  |
| pMG006 | pBJ114 todK_D276A_ | oPH972 | A | above |  | DZ2 | pBJ114 |
|  |  | oPH1634 | B | GCGGATCGCATGGGTGACGATGCCCATG |  |  |  |
|  |  | oPH1635 | C | CACCCATGCGATCCGCAGCCCGCTGGGC |  |  |  |
|  |  | oPH975 | D | above |  |  |  |
| PH2011 |  | oPH1636 | E_wt_ | TGGGCATCGTCACCCATGAC |  | todK_D276A_ loop-out clones | N/A |
|  |  | oPH1637 | E_mut_ | TGGGCATCGTCACCCATGCG |  |  |  |
|  |  | oPH994 | F | above |  |  |  |
| pMG007 | pBJ114 todK_D279A_ | oPH972 | A | above |  | DZ2 | pBJ114 |
|  |  | oPH1638 | B | CAGCGGCGCGCGGATGTCATGGGTGAC |  |  |  |
|  |  | oPH1639 | C | ATCCGCGCGCCGCTGGGCGCCATCATG |  |  |  |
|  |  | oPH975 | D | above |  |  |  |
| PH2012 |  | oPH1640 | E_wt_ | GTCACCCATGACATCCGCAGC |  | todK_D276A_ loop-out clones | N/A |
|  |  | oPH1641 | E_mut_ | GTCACCCATGACATCCGCGCG |  |  |  |
|  |  | oPH994 | F | above |  |  |  |
| pMG013 | P*_pilA_*-*todK* | oPH944 | A | AT**GGATCC**GCTGATCGACAGTTATCGTC | BamHI | DZ2 | pFM18 |
|  |  | oPH1670 | B | GGTGGGGGGCATGGGGGTCCTCAGAGAAGG |  |  |  |
|  |  | oPH1671 | C | ACCCCCATGCCCCCCACCCCCGCCAAGAAGC |  |  |  |
|  |  | oPH1672 | D | CAT**AAGCTT**TTAGTCGCGCGGGCTTCC | HindIII |  |  |
| pMG014 | P*_pilA_*-*todK*_H275A_ | oPH944 | A | AT**GGATCC**GCTGATCGACAGTTATCGTC | BamHI | PH2010 | pFM18 |
|  |  | oPH1670 | B | GGTGGGGGGCATGGGGGTCCTCAGAGAAGG |  |  |  |
|  |  | oPH1671 | C | ACCCCCATGCCCCCCACCCCCGCCAAGAAGC |  |  |  |
|  |  | oPH1672 | D | CAT**AAGCTT**TTAGTCGCGCGGGCTTCC | HindIII |  |  |
| pMG015 | P_pilA_-*todK*_D276A_ | oPH944 | A | AT**GGATCC**GCTGATCGACAGTTATCGTC | BamHI | PH2011 | pFM18 |
|  |  | oPH1670 | B | GGTGGGGGGCATGGGGGTCCTCAGAGAAGG |  |  |  |
|  |  | oPH1671 | C | ACCCCCATGCCCCCCACCCCCGCCAAGAAGC |  |  |  |
|  |  | oPH1672 | D | CAT**AAGCTT**TTAGTCGCGCGGGCTTCC | HindIII |  |  |
| pMG016 | P*_pilA_*-*todK*_S279A_ | oPH944 | A | AT**GGATCC**GCTGATCGACAGTTATCGTC | BamHI | PH2012 | pFM18 |
|  |  | oPH1670 | B | GGTGGGGGGCATGGGGGTCCTCAGAGAAGG |  |  |  |
|  |  | oPH1671 | C | ACCCCCATGCCCCCCACCCCCGCCAAGAAGC |  |  |  |
|  |  | oPH1672 | D | CAT**AAGCTT**TTAGTCGCGCGGGCTTCC | HindIII |  |  |
| pMG018 | TodK protein overproduction | oPH601 | forward | GAC**GAATTC**ATGCCCCCCACCCCCGCC | EcoRI | DZ2 | pET32a |
|  |  | oPH604 | reverse | GCG**GTCGAC**TTAGTCGCGCGGGTTCC | SalI |  |  |
| pMG019 | TodK_H257A_ protein overproduction | oPH601 | forward | GAC**GAATTC**ATGCCCCCCACCCCCGCC | EcoRI | PH2010 | pET32a |
|  |  | oPH604 | reverse | GCG**GTCGAC**TTAGTCGCGCGGGTTCC | SalI |  |  |
| *-* | *csgA* qPCR | oPH378 | forward | GGGCGATACCGTCGAAGC |  | *-* | *-* |
|  |  | oPH379 | reverse | CTGTCGTCGTCTCCCACGTC |  |  |  |
| *-* | *fruA* qPCR | oPH252 | forward | CGTCACGGAAGGCATCAATC |  | *-* | *-* |
|  |  | oPH253 | reverse | CGAGATGATTTCCGGTGTGC |  |  |  |
| *-* | *mrpC* qPCR | oPH353 | forward | GGAGGCCATCGACTTCAAGG |  | *-* | *-* |
|  |  | oPH354 | reverse | GGCCGGACTTCAGCAGGTAG |  |  |  |
| *-* | *espA* qPCR | oPH369 | forward | CGACGTTGGATGAACTCACG |  | *-* | *-* |
|  |  | oPH370 | reverse | GCACGGTGACGTCGGAAC |  |  |  |
| *-* | 16S qPCR | oPH235 | forward | AACTGTTGTGCTTGAGTGCCG |  | *-* | *-* |
|  |  | oPH236 | reverse | ATCTAATCCTGTTTGCTCCCCAC |  |  |  |

^a^ Description of insertion plasmids

^b^ Primers listed as A, B, C, D were used for generating mutations via overlap PCR and primers listed as E and F were used for screen of point mutations as described in detail in (Lee et al., 2010). For direct amplification of target DNA, primers are listed as forward and reverse. Restriction site sequences are in bold. Mutated bases are underlined.

**Supplemental Methods**

*Table S2. Parameters for the NLA image-analysis pipelines*.

| Pipeline | Parameter^a^ | Value^b^ | Script |
| --- | --- | --- | --- |
| Radial darkness (Fig. 1C, S1A,B, S2A,B) | Spatial scale | 0.005123 mm px⁻¹ (5.123 µm px⁻¹) at full resolution | cf_wt_dtodk_n6.py |
|  | Working resolution | half-resolution (2× averaging downscale) |  |
|  | Local-background Gaussian σ | ≈100 full-res px (≈0.5 mm) |  |
|  | Radial smoothing window | 45 full-res px |  |
|  | Centre and growth-front radius | set by hand in the interactive circle editor; used verbatim |  |
|  | Coverage rule | solid where the sampling ring is ≥95% within frame; dotted below |  |
| Granularity / texture  (Fig. 1E, S1D, S2C) | Working resolution | half-resolution (2× averaging downscale) | granularity script |
|  | Illumination correction | divide by Gaussian blur (σ ≈ 120 full-res px), rescaled to background mean |  |
|  | Band-pass | difference-of-Gaussians, σ ≈ 2 minus σ ≈ 12 full-res px |  |
|  | Metric | population standard deviation of the band-passed image within the ROI (arbitrary units) |  |
|  | Region of interest | fixed 720 px full-res radius; whole 0–1.0 R, core 0–0.65 R, ring 0.65–1.0 R |  |
|  | Value scaling | none (raw SD, 3 dp); wild-type normalisation applied downstream to per-strain means |  |
| Body detection and spatial statistics  (Fig. 1D, S1C, Table 1) | Detector input | local-background darkness map (not the raw image) | cf_nnd_fullrun.py |
|  | LoG blob detection | min σ = 4, max σ = 12, 8 scales, threshold 0.05 (on normalised darkness map) |  |
|  | Darkness gate | peak ≥ 13 a.u. and mean ≥ 5 a.u.; candidates < 10 px radius additionally require mean ≥ 9 or peak ≥ 20 a.u. |  |
|  | Merge rule | largest-first; absorb a smaller candidate overlapping a retained one by > 1/6 of the smaller candidate’s area |  |
|  | Spatial statistics | count; density (bodies mm⁻²); mean nearest-neighbour distance (mm) |  |
|  | Validation | detections eye-validated against the source images |  |

^a^ Parameters correspond to the analysis scripts archived at Zenodo (DOI: 10.5281/zenodo.21538883). LoG, Laplacian-of-Gaussian

^b^Values are given as full-resolution equivalents unless noted; pipelines operating at half resolution are indicated. a.u., arbitrary units; NND, nearest-neighbour distance; ROI, region of interest

**Detailed NLA image analysis methods**

***Validation approach.*** Because no ground-truth annotation existed for these images, each component of the pipeline was validated against expert visual assessment of the source images: candidate detections were overlaid on the images and inspected, and radial profiles were drawn on the spots and checked. Parameter values were fixed by this visual validation and then applied uniformly to every image; no parameter was tuned per image, apart from the interactive spot-center and growth-front settings described below.

***Image handling and calibration.*** Source images were 2048 × 1536 px RGB brightfield stereomicrographs, one field per developmental spot, converted to grayscale (luminance-weighted) before any measurement. The spatial scale was 0.005123 mm/px (5.123 µm/px) at full resolution. The texture and darkness pipelines operated at half resolution (1024 × 768 px) for efficiency; lengths below are given as full-resolution equivalents. Downscaling used averaging (area interpolation) rather than stride sampling, which is not equivalent.

***Texture (granularity) metric.*** Aggregation was quantified as image granularity, defined as the standard deviation of a band-pass-filtered, flat-field-corrected image within a fixed circular region. For each image, the grayscale frame was downscaled twofold by averaging. Illumination was corrected by dividing the image by a strongly blurred copy of itself (Gaussian σ ≈ 120 full-resolution px) rescaled to the mean background level, removing the smooth illumination gradient while preserving structure. A difference-of-Gaussians band-pass (σ ≈ 2 minus σ ≈ 12 full-resolution px) isolated structure at the fruiting-body scale, and the metric was the population standard deviation of this band-passed image within the region of interest. Because the operation is a band-pass, the metric is insensitive both to the illumination gradient and to absolute image brightness, responding only to structure at the scale of interest. The region of interest was a fixed circle of 720 full-resolution px radius, identical for every spot (the spots were equal-volume drops of similar size), centered on an automatically fitted drop-edge circle: a texture-based colony mask was fitted by algebraic (least-squares) circle fitting with iterative outlier rejection, and only the fitted center was used, with the region radius held fixed. Three regions were reported: whole spot (0–1.0 R), inner core (0–0.65 R), and outer ring (0.65–1.0 R, the spot periphery). Values were raw standard deviations in arbitrary units; normalization to the same-experiment wild-type mean was performed separately on per-strain means and was not applied to the stored values.

***Darkness metric.*** Fruiting-body maturation and spatial organization were quantified from a local-background-referenced darkness map. Darkness was defined as clip(Gσ − I, 0), where I is the grayscale image and Gσ is a Gaussian-blurred estimate of the local background (σ ≈ 100 full-resolution px, ≈ 0.5 mm). Subtracting the image from its local-background estimate and clipping at zero yields a positive value wherever the image is darker than its immediate neighborhood. Because the reference is local (≈ 0.5 mm) rather than global, spatially broad features — bare agar, the illumination gradient, and the underlying vegetative-cell lawn — are absorbed into the background and removed, so that only compact structures dark relative to their surroundings survive. The metric therefore reports each body's darkness in excess of the local vegetative and agar background rather than absolute pigmentation. This behavior was confirmed by the non-developing Δ*mrpC* control (Fig. S1A,B), which is rich in spotted cells but devoid of fruiting bodies and read flat at approximately 2 a.u. against a wild-type peak of approximately 13.5 a.u.

***Radial darkness profile.*** For spatial-organization analysis, the profile was the azimuthal (360°) mean darkness in 1-px annuli measured outward from the spot center, smoothed with a 45-px moving average, and plotted against absolute radial distance (mm). Because it is an azimuthal mean, the profile reports a local areal darkness density at each radius rather than a body count. For the 48-h radial analysis, the spot center and growth-front radius were set by hand for each image using a purpose-built browser-based circle editor and used verbatim without automatic refitting; this human-in-the-loop step was necessary because smooth wild-type colony fronts defeat automatic edge detection, and an incorrect front radius displaces the analysis windows. Where automatic front detection was used, it was validated against the hand-set circles at 48 h (detected/hand-set radius ratio 0.967 ± 0.064, n = 37). The fraction of each annulus lying within the image frame (coverage) was tracked; profiles were drawn as solid lines where coverage was complete and dotted beyond. Profiles represent the mean of six independent biological replicates, with the shaded band showing the standard error of the mean, and are presented descriptively; no hypothesis test was applied to the curves.

***Fruiting-body detection.*** Fruiting bodies were detected on the local-background darkness map (not the raw image), which makes detection robust to the illumination gradient and the vegetative lawn. Candidate bodies were found by Laplacian-of-Gaussian (LoG) blob detection over a range of scales (scikit-image; van der Walt et al., 2014; minimum σ = 4, maximum σ = 12, 8 scales, detection threshold 0.05 on the normalized darkness map). Candidates failing an absolute darkness floor (peak < 13 a.u. or mean < 5 a.u.; small candidates of radius < 10 px additionally required mean ≥ 9 or peak ≥ 20 a.u.) were rejected, preventing spurious detections in near-agar regions. Overlapping candidates were resolved largest-first: a smaller candidate overlapping a retained larger one by more than one-sixth of the smaller candidate's area was merged into it. This threshold was set by visual validation — loose enough to fuse the multiple detections attracted by a single elongated or irregular body, yet tight enough to preserve genuinely adjacent bodies in dense Δ*todK* spots. The detector returned, per body, a centre and a fitted circle radius; the radius is a relative size proxy, not a calibrated measurement. Detections were validated by overlay inspection (Fig. S1C). A small number of elongated bodies still split into two detections, and the densest Δ*todK* spots are the hardest to resolve, so Δ*todK* counts are, if anything, under-called — which makes the reported count difference conservative. Wild-type bodies migrate and leave elongated smears, which were treated as single bodies.

***Spatial statistics.*** From the detected body centers, three statistics were computed per spot: count (number of bodies within the spot region); density (count per mm² of spot area); and mean nearest-neighbor distance (NND; for each body, the Euclidean distance to the nearest other center, averaged over bodies and converted to mm). Statistical significance for the wild-type vs Δ*todK* comparison was assessed by a two-sided Mann–Whitney U test on the six per-experiment means, chosen because the two strains had markedly unequal variance.
